# Sedentary life accelerates epigenetic ageing in King penguins

**DOI:** 10.1101/2024.09.24.614416

**Authors:** Robin Cristofari, Leyla Davis, Gaël Bardon, Flávia A. Nitta Fernandes, Maria Elena Figueroa, Sören Franzenburg, Michel Gauthier-Clerc, Francesco Grande, Richard Heidrich, Mikaela Hukkanen, Yvon Le Maho, Miina Ollikainen, Elodie Paciello, Patrick Rampal, Nils C Stenseth, Emiliano Trucchi, Sandrine Zahn, Céline Le Bohec, Britta S. Meyer

## Abstract

Advances in medicine and food security have contributed to an increase in human lifespan^1^. Yet, the associated rise in sedentary behaviour and in obesity^2,3^ already threatens these gains^4^. Indeed, a growing body of evidence supports the central role of nutrient sensing and energy management pathways in regulating ageing rate and healthspan^5,6^, but the diversity of human lifestyles challenges our ability to identify the genetic and epigenetic drivers of this age acceleration. Here, we examine how the transition of wild King penguins to zoo husbandry can closely mimic the shift to a Western lifestyle in humans, and shed light on evolutionarily conserved epigenetic changes in responses to sedentary conditions. We show that, just like modern humans, zoo-housed King penguins experience an extended lifespan, but this comes at the cost of accelerated epigenetic ageing throughout life. This accelerated ageing is associated with differential methylation in key growth and maintenance pathways including the mTOR and PI3K/Akt networks, as well as in specific pathways of lipid-rich diet adaptation and heart-function. Our results demonstrate the deeply conserved link between sedentary behaviour and food availability on the one hand, and age acceleration on the other. Such evolutionary evidence may in turn help us to improve risk detection and, ultimately, therapeutics for lifestyle-induced age acceleration in humans^7^.

## Main text

At the epidemiological level, there is a clear association between sedentary behaviour, obesity, and accelerated ageing phenotypes in humans8,9. Clinically, this association is reflected in the convergence of processes that include the dysregulation of nutrient sensing, metabolic disruption, impaired mitochondrial function, and a chronic inflammatory state associated with cellular senescence8,10. Within the nucleus, this translates into telomere shortening, associated with increased epigenetic drift11,12, accelerated clock-like epigenetic changes11,13, and decreased genome stability8. In short, sedentary behaviour and obesity accentuate most, if not all, of the hallmarks of ageing14,15, and in humans, this is directly reflected in epigenetic age acceleration (EAA)11,13,16. Thus, the resulting effect on ageing seems primarily pathological, stemming from a general derangement of homeostasis8. Ultimately, this means that an individual’s lifelong history of nutrient intake and energy management, although difficult to estimate in humans, is likely to be a key driver of EAA.

These findings have naturally led to the idea that improving nutrient intake and physical activity might be a shortcut to lifespan extension. Indeed, calorie restriction (CR)12,17–19, physical activity20, as well as direct manipulation of nutrient-sensing pathways21,22, have shown positive effects on ageing rates. However, it is still unclear whether such benefits add up over a lifetime23: the spectacular benefits of CR in the laboratory often disappear in a natural environment24, while its “hidden costs” become more apparent25. Studies conducted in mice are limited by the species’ short lifespan and characteristic metabolism26, making them a limited model for examining some mechanisms in humans27. Clinical trials are scarce28, and reliable long-term data dietary intake in humans extremely difficult to obtain29, especially as modern food sources - in particular ultra-processed foods - are increasingly thought to act on ageing independently of their calorie content30. As a result, there is still fierce debate about whether the health span benefits observed in the laboratory also apply to natural, lifelong, periodic CR in a long-lived species such as humans27,28,31.

### Penguins as a model for the Western lifestyle

Here, we test the effects of lifelong manipulation of physical activity and food intake on ageing rate using a novel model, the king penguin (Aptenodytes patagonicus). Aptenodytes penguins have long been studied for their outstanding fasting abilities, and are able to go without food for up to eight weeks - a fast that, despite its extreme length, shares key physiological traits with human fasting32. As part of their breeding cycle, King penguins alternate between prolonged CR and intense physical activity, with foraging trips requiring them to swim up to 1,200 km in the open ocean in a few days33; and unlike any other model to date, penguins voluntarily stop eating in the wild, removing one of the current barriers standing between laboratory and real-life conditions24,25.

For larger, long-lived wild animals such as king penguins, the transition to zoo husbandry has relevant similarities to modern Western human lifestyle. Protection from predators, reliable food sources and advanced veterinary care reduce extrinsic mortality hazards, and can increase average lifespan34. On the other hand, despite best husbandry practices, physical activity is necessarily reduced. Indoors housing is often required, entailing artificial lighting and simulated seasonality. While these conditions are theoretically beneficial overall for longevity, depression-like emotional states can follow35, and overweight is a recurrent issue in zoo-housed animals, often requiring specific dietary management36. For a species such as the king penguin, this creates a unique experimental system in which CR and high energy expenditure are effectively a control state, and sedentary behaviour and continuous feeding are an experimental manipulation.

Specifically, we hypothesise that introducing a Western-human-like evolutionary mismatch by inducing lifelong sedentary behaviour and increased food availability would lead to accelerated ageing in zoo-born king penguins. A candidate mechanism for this is the chronic activation of pro-growth pathways, and to the associated suppression of cellular repair and autophagy mechanisms, e.g. via counter-acting properties of the mTOR pathway and other mechanisms balancing growth and somatic maintenance. If observed, such regulatory changes associated with EAA would independently validate the currently hypothesised impact of lifestyle changes on ageing rate, and would provide novel evidence for a robust, evolutionarily-conserved link between nutrient regulation, physical activity and ageing.

### Epigenetic age acceleration in zoo-housed penguins

To estimate biological ageing in king penguins, we relied on a CpG methylation-based epigenetic clock, as this approach has been effective at predicting frailty and mortality risk in a wide range of species^37,38^. In order to calculate EAA in king penguins in the absence of an externally-validated epigenetic clock, we produced whole-genome sequencing data for 64 known-age samples for male King penguins, 34 of which from the wild (Possession Island, Crozet archipelago, in the Indian sector of the Southern Ocean), and 30 born and housed at the zoo (10 at Zoo Zurich, Switzerland, and 20 at Loro Parque, Tenerife, Spain). After read processing, we obtained whole-genome CpG methylation data at an average median depth of 29.5X across samples (range: 20X to 41X). After pooling methylation calls for both strands, our filtered dataset contained a total of ∼10,000,000 high-quality CpG sites (see Supplementary Table S3). We did not find any evidence for genetic divergence between the wild and zoo groups (see Supplementary Materials S3), in keeping with the small number of generations elapsed since introduction from the wild (7 generations at most in our sample). We therefore exclude a systematic genetic basis for EAA in our experimental setup.

To infer epigenetic age, we used the residuals from an elastic net regression fitted to the principal components of variance in CpG methylation levels^38,39^. The final inter-group difference was highly significant when modelling predicted epigenetic age against calendar age: correcting for library conversion efficiency and genome-wide average CpG methylation level, the effect of living condition on average EAA at the zoo is estimated as 6.48 years (SE = 0.74, p-value < 0.0001). When allowing for separate slopes, however, the difference in slope was not significantly different from zero (−0.12, SE = 0.08, p-value = 0.16). Our final model therefore included only inter-group difference in intercept (baseline age acceleration), and had an overall excellent fit (RMSE = 2.48 years, R^2^ = 0.91) (Fig. 1A and 1B). In that model, EAA was not different between individuals from the two different zoos included in the study, demonstrating that the difference observed in EAA is driven by lifestyle rather than the population sampled (t-statistic = −0.15, p-value = 0.88) (see Methods and Supplementary materials S4 for further validation of this result).

**Fig. 1.**
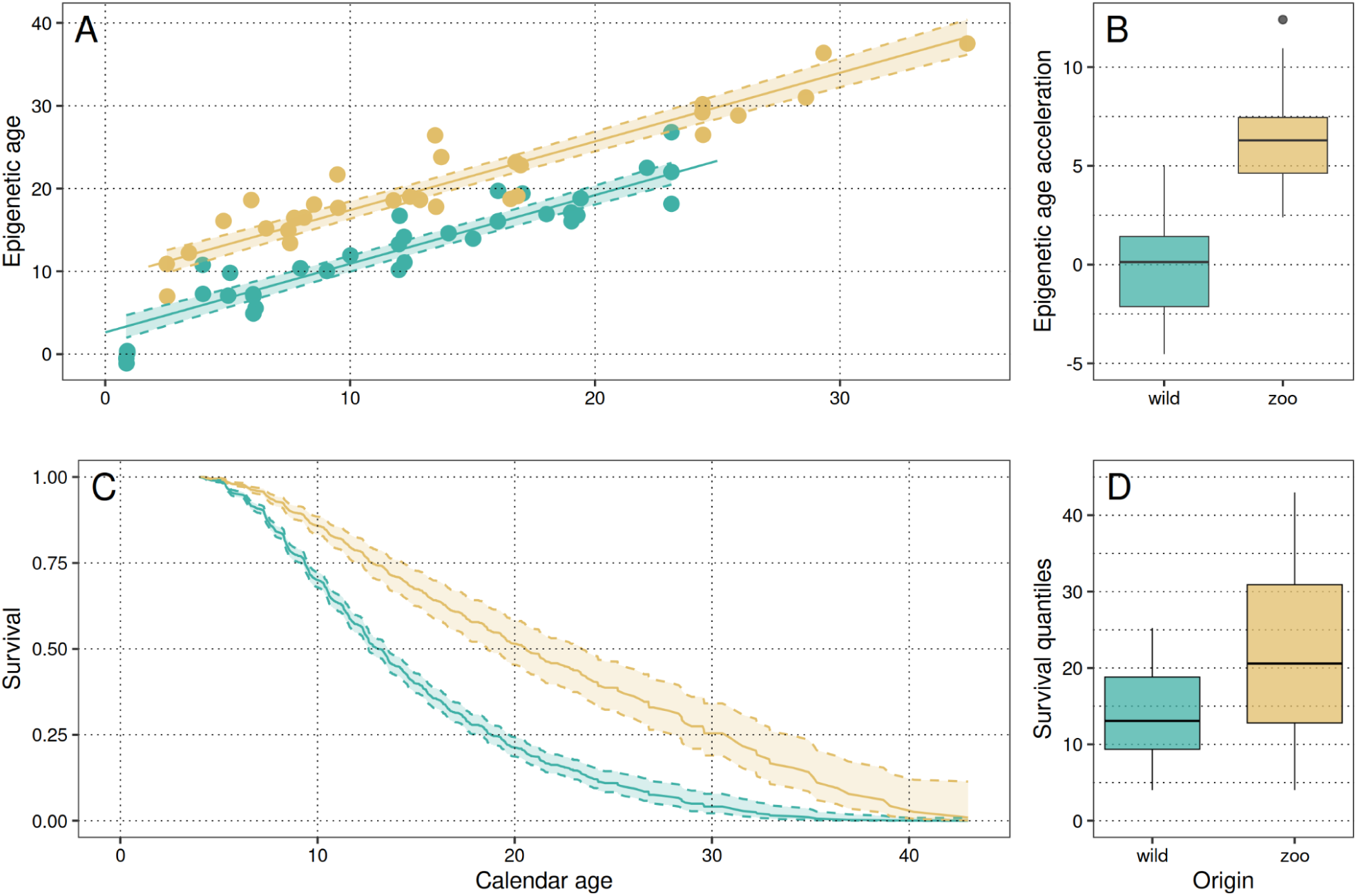
| Epigenetic age acceleration and survival in wild and zoo-housed King penguins. In green, wild birds, in tan, zoo birds; boxplots provide color reference. (A) Comparison of epigenetic age versus calendar age between wild and zoo-housed King penguins, with 95% confidence intervals. Data points represent raw age predictions derived from CpG methylation levels. (B) Distribution of age acceleration in wild and zoo-housed King penguins. (C) Cox proportional hazard model illustrating survival probabilities for wild and zoo-housed male King penguins. (D) Survival quantiles from the Cox-PH model (median and quartiles).

As an additional control, we applied our approach to a human test dataset of similar characteristics (see Methods below), but using as a contrast the strongest single known behavioral factor affecting EAA in humans - smoking^40^. Our approach for EAA in smokers versus non-smokers yielded results that were highly consistent with the independently-trained PCPhenoAge and PCGrimAge clocks^39^ (see Supplementary material S4 and Fig. S5). These results strongly support the view that king penguins housed in zoos display higher levels of EAA than their wild counterparts: relative to the species longevity (∼40 years), this difference in EAA is comparable in magnitude to the extremes observed in humans, here between smokers and non-smokers, and make the zoo transition a major driver of EAA.

### Zoo-housed penguins have increased survival

In contrast with the observed age acceleration in zoo conditions, survival analysis using Cox proportional-hazard modelling showed that zoo-housed penguins lived longer on average than their wild counterparts. This analysis included 1,895 wild and 305 zoo-housed king penguins, with nearly no differences between sexes (Fig. 1C and 1D). For zoo-housed birds, the median survival age (with 95% confidence intervals) was 20.7 years [18.7-23.6] for males and 20.8 [18.8-23.7] for females. In comparison, the median survival age for wild birds was significantly lower, at 13.5 years [13.0-14.1] for both sexes. The hazard ratio comparing zoo-housed to wild conditions was 0.46 [0.38-0.55], indicating a substantial advantage for penguins housed in zoos (the methodology and additional analyses can be found in the Methods section and Supplementary Material S2). Age-specific survival is lower at younger ages in the wild, reflecting the strong effect of extrinsic mortality (predation and starvation at sea in particular), while protection from accidental death, food security, and a high level of medical care shelters zoo-housed penguins until more advanced ages - an observation consistent with previous results in zoo-housed mammals^34^. As a result, at equal age and frailty level, survival is considerably increased at the zoo compared to the wild.

We examined whether selective disappearance was a likely cause of this difference, with the proportion of frail and age-accelerated individuals decreasing more rapidly through time in the wild. However, the absence of correlation between EAA and age either in the wild (Spearman’s rank test p-value = 0.28), in the zoo (0.13), or in both (0.52) strongly suggests that selective disappearance is not a major determinant of EAA in either condition, as strong survival-bias would induce a change in age-class individual composition. Yet, all the individuals included in this study are still alive, so that direct association between age acceleration and lifespan is, at this stage, impossible. Overall, we suggest that birds housed in zoo conditions undergo accelerated ageing, but that this age acceleration is compensated by their highly protected lifestyle, allowing for survival to advanced ages in conditions of frailty, which would rule out survival in the wild.

### Differentially methylated genes

In order to understand the determinants of age acceleration at the zoo, we searched for differentially methylated regions (DMRs) between zoo-housed and wild birds independently of age-acceleration signal. We beta-corrected individual CpG methylation levels for chronological age, EAA and genome-wide methylation level (see Methods), in order to only retain CpG methylation signal that is consistently different between groups, but does not covary with either chronological age, or with EAA. DMRs inferred from this signal should thus reflect baseline “methylation switches” that oppose zoo and wild penguins in a time- and EAA-independent manner. This approach identified a total of 600 candidate DMRs (Fig. 2A), clustered in or near 292 known genes, and 36 predicted open reading frames (ORFs) in the king penguin - a result well outside random expectations (see Supplementary Material S5 and Supplementary Fig. S9).

### Overrepresentation of growth and maintenance pathways

Pathway overrepresentation, tested against the Reactome pathway database^41^, clustered these genes into 11 FDR-corrected “super-paths” (See Supplementary Table S5 for the full results). Our gene set was significantly enriched for pathways centrally involved in cell growth, and in the coupling of nutrient sensing to ageing and age acceleration^5,6,44^. These include **(1) the MAPK and RAS-RAF-MEK-ERK pathways** (18 genes, FDR-corrected p-value = 0.0003), **(2) the PI3K/Akt network** (7 genes, 0.0013), **(3) Notch signaling** (4 genes, 0.0012), **(4) ALK signaling** (5 genes, 0.0049), **(5) EGFR signaling** (5 genes, 0.0008), and **(6) the Rho GTPase Ras subfamily** (4 genes, 0.0042). These pathways are consistently involved in **(a) the regulation of cell growth** (including mTORC2 subunit MAPKAP1, mTOR regulator SIK3, the platelet-derived growth factor receptor A PDGFRA, or the fibroblast growth factor receptor 3 FGFR3^42,43^), **(b) apoptosis** (including p53 effector PERP, proto-oncogene REL, apoptosis-inducing factor mitochondria-associated AIFM3, and both EVA1B and EVA1C, the two known paralogs of the regulator of programmed cell death EVA1A), and **(c) DNA-damage response** (e.g. HSF1, ZGRF1, BUB1, SSBP2, SLX4, GADD45A, WRD76 or USP28) – see Fig. 2B for a representative subset of these genes, and Supplementary Table S4 for the full list of result. The clear overrepresentation of central growth and maintenance pathways in the EAA-independent differential methylation between zoo and wild conditions strongly supports our hypothesis, and suggests that the induction of a sedentary, well-fed lifestyle has a direct influence on core metabolic regulation in king penguins.

**Fig. 2.**
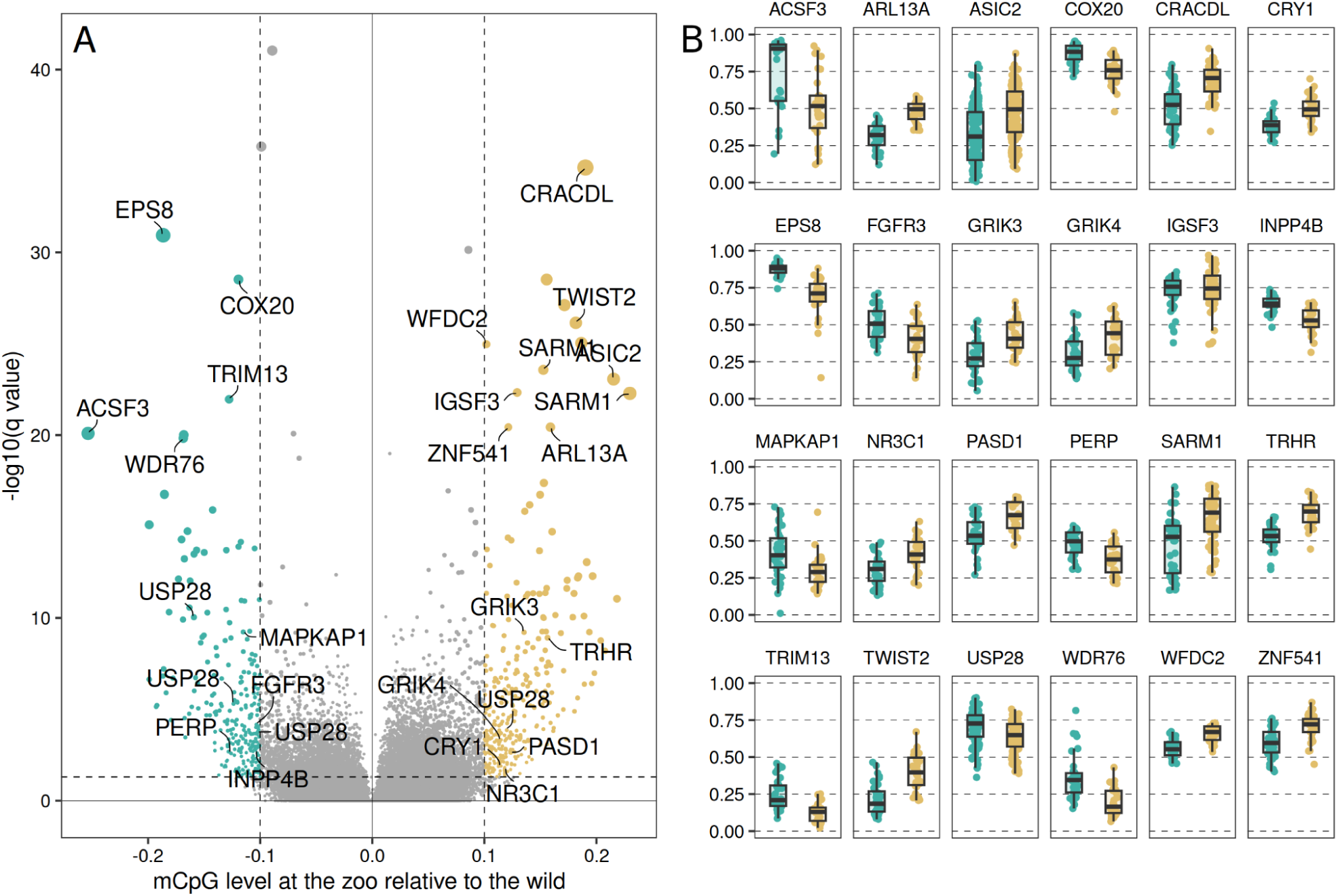
| Age-independent differentially methylated regions (DMRs) between zoo and wild King penguins. (A) Volcano plot showing all DMRs. X axis: relative change in methylation level (0: no difference, positive, in tan: hypermethylated at the zoo, negative, in green: hypomethylated at the zoo). Y axis: −log10 of the Benjamini-Hochberg-corrected q-value based on Mann-Whitney U test. Coloured areas: DMRs retained as significant. Labels show top DMRs as well as some chosen relevant DMRs discussed in the main text. The full list is provided in Supplementary Table S4. (B) Adjusted methylation level at the DMRs labelled in the volcano plot. In green (left), wild birds, in tan, zoo birds; boxplots provide color reference. Methylation levels are beta-corrected for age, EAA and genome-wide mean CpG methylation.

### Change in diet, heart function and physical activity

Our results include DMRs associated with genes directly involved in coping with excessive nutrient intake, including INPP4B^45^ and ASIC2^46^ and NAD hydrolase SARM1 - the latter centrally involved in cellular response to glucose, growth and apoptosis, whose deletion specifically improves function in the diabetic heart^47^ and mitigates the effects of high-fat diets^48^. They also functionally overlap genes recently associated with heart function and physical exercise in genome-wide association study (GWAS) in a large human cohort^49^, including GRIK3 and GRIK4 (the first subunit of this receptor, GRIK1, is linked to heart rate increase and recovery in humans^50^). These overlaps also include RNF220, KCNH5 (its paralog KCNH8 is a GWAS in humans), and CAVIN-1 (a main interactor of CAV1 identified as a GWAS). It is remarkable that these epigenetic changes can be reliably detected in blood, as opposed to tissues more centrally involved in diet and physical activity (e.g. muscle, liver or adpipose tissue). This underlines the value of a non-invasive matrix such as peripheral blood for insights on whole-organism methylome^51^. These results primarily suggest that (1) zoo-housed King penguins may mobilise a wide array of genes across tissues to specifically compensate for a change in diet lipid composition and/or mitigate the negative effects of food abundance and (2) the suppression of intense physical activity in a species adapted to sustained efforts, that can routinely swim several hundred to thousands kilometers in a few days^33^, contributes directly to systemic epigenetic change. Further research on these genes may provide candidates for mitigating the effects of adiposity and sedentary behaviour in humans.

### Effects of life indoors

At the zoo, birds have a significantly hypermethylated DMR in the promoter of NR3C1, the glucocorticoid receptor (GCR), and ∼3kb upstream of TRHR, the thyrotropin-releasing hormone receptor. In humans, increased NR3C1 promoter methylation is associated with decreased GCR expression and psychosocial stress^52^; and it correlates negatively with circulating corticosterone levels in seagulls^53^. This suggests low situational stress and low nutritional stress (CG levels increase with CR in King penguins^32^). Hypermethylation upstream of TRHR, on the other hand, may be associated with lowered TRHR expression and a blunted thyroid-stimulating hormone (TSH) response to TRH, a trait regularly observed in depression in humans. The observed methylation patterns are thus consistent with a sheltered lifestyle with lower GC-mediated stress. Characteristically, the circadian clock pathway is also significantly enriched in our DMR results (5 genes, FDR-corrected p-value = 0.0039, including cryptochrome circadian regulator CRY1 and in PASD1, a known clock repressor^54^). These results strongly suggest that sedentary lifestyle has a broad impact on zoo-housed king penguin physiology. A possible association with a depression-like state remains speculative and should be investigated further - but acute stress is unlikely to be causal in this case.

## Discussion

Our results bring novel, independent evidence to support the idea that the transition to a sedentary, sheltered and food-stable lifestyle is a conserved cause of age acceleration across a wide evolutionary divide. Importantly, they suggest that EAA is driven by the suppression of periodic CR and of physical activity, rather by than lipotoxicity or other direct pathological effects of overweight: indeed, penguins in this study were not clinically obese (at the zoo, inter-breeding body mass was 11.8 kg, sd = 2.2, not markedly different from a ∼12 kg inter-breeding wild bird^32^). Instead, differential methylation analysis points to pathways deeply involved in the systemic regulation of metabolism, growth and maintenance, rather than at specific effects on e.g. obesity-induced inflammation. These results considerably strengthen the hypothesis that evolved metabolism mismatch, rather than side-effects of the associated cultural changes, is at the root of the ongoing increase in age-acceleration diseases in humans.

We also underline the fact that the observed EAA does not appear to be the result of an overall pace-of-life acceleration, as breeding output at the zoo is also decreased compared to the wild^55^. This decoupling of age acceleration and pace-of-life suggests the existence of a large, unexploited margin for lifestyle-based lifespan extension: in the penguin model, a “best of both worlds” lifestyle combining the physical activity, native diet and slower ageing rate of wild birds, together with the reduction of extrinsic mortality and medical care of the zoo would, in theory, result in a 20% increase in survival. In humans, this would be equivalent to a ∼12 to 15 years gain in life expectancy - more than the estimated years of life lost to severe obesity in the Western lifestyle^56^. While such an extrapolation from penguins to humans is speculative at best, it underlines the magnitude of the gains in life expectancy that could possibly be achieved by acting solely on lifestyle factors - in modern humans as well as in animals under their care.

## Methods

### Sample collection

DNA was collected from the whole peripheral blood of 64 male King penguins of known age (see Supplementary material S0 for details). The free-ranging individuals (N = 34) were followed since fledging in the wild (Possession Island, Crozet Archipelago), as part of a long-term monitoring project^57^, whereas the individuals from zoos (N = 30) were hatched in European zoos, and housed at Zoo Zurich, Switzerland (N = 10), and Loro Parque, Spain (N = 20), at the time of sampling. In the wild, the sampling was approved by the French ethics committee (latest: APAFIS#29338-2020070210516365) and the French Polar Environment Committee, and permits to handle the animals and access the breeding sites were issued by the “Terres Australes et Antarctiques Françaises”. In zoos, samples were collected as part of routine veterinary examination. The choice to restrict the analysis to males only was guided by the asymmetry in fasting behaviours between males and females in this species: males systematically take the first turn in incubating the egg, fasting for at least 6 weeks on average at the start of reproduction, while females go foraging at sea immediately after egg laying, staying on land for ∼ 3 weeks^58^. Males thus offered a higher contrast in fasting behaviour between wild and zoo conditions.

### Sequencing and data processing

Whole-genome enzymatic-conversion methylome sequencing libraries were prepared using the NEB EM-seq kit^59^, and sequenced on 6 sequencing lanes on the Illumina NovaSeq 6000 platform at CCGA in Kiel, Germany. Raw reads were trimmed using trimmomatic^60^, removing leading and trailing bases if the quality score dropped below 3 and retaining only reads of length > 36 bp. We used BCREval^61^ to assess conversion efficiency during library preparation based on false-positive non-CpG methylation calls in telomeric repeats. Reads were aligned to the King penguin’s reference genome^62^ using bsbolt^63^ (v1.5.0), and deduplicated and filtered using samtools^64^ (v1.16.1). Individual methylated cytosines were called using bsbolt. In vertebrates, DNMT1 activity ensures that CpG methylation is normally symmetric across strands^65^: therefore, we calculated per-CpG average cytosine methylation level by pooling C and T calls from both strands at each CpG dinucleotide^66^. We masked from our dataset all known C/T (or G/A) SNP positions based on a curated database of 40 King penguin whole genomes (Fernandes et al. *in prep*). Mean conversion efficiency as assessed through telomeric methylcytosine false-positive call rate was high at 99.5% (range: 96.6% to 99.9%): this method yields results that are very close to the internal PhiX control, but that reflect more directly the conversion efficiency on the target species’ DNA^61^.

### Data annotation

The genome reference used^62^ was re-annotated for this study. Briefly, we assessed assembly completeness using BUSCO^67^. It was found highly complete (C:95.6% [S:95.0%, D:0.6%], F:1.5%, M:2.9%, n:8338), although the published annotation included a large proportion of incomplete gene models. We used BRAKER^68^ to generate genome annotations based on (i) whole-transcriptome data for the wild King penguin, aligned to the King penguin genome using HiSAT2^69^, and (ii) the OrthoDB^70^ vertebrate protein reference dataset. For additional details on the transcriptome data used, see Paris et al. in preprint^71^. Untranslated regions (5’- and 3’-UTRs) were annotated using GUSHR (https://github.com/Gaius-Augustus/GUSHR), based on the same sources of information. Cis-regulatory promoter elements were defined either as the 1500bp upstream, and 500 bp downstream of the transcription start site (TSS)^72^. CpG islands (CpGi) were defined as regions of high CpG density using CpGCluster^73^. Gene annotations were established using protein-protein BLAST^74^ against avian proteins.

### Epigenetic age acceleration (EAA)

In order to assess epigenetic age acceleration in wild and zoo individuals, we followed the approach outlined by Higgins-Chen and colleagues^39^, performing first a principal component analysis using MethylKit^75^. However, we reduced the extremely large starting CpG set in two steps: (*i*) we retained only sites with an average single-stranded depth of 10 to 25x, and an individual sample depth of 5 to 50x, no missing data in any individual, and a standard deviation in CpG methylation level of at least 0.1 (2,670,062 CpGs), and (*ii*) to retain only broadly informative sites, we calculated Pearson’s correlation coefficient between individual age and CpG methylation level at each CpG, and only retained sites with an R^2^ ≥ 0.2 and a p-value ≤ 0.05 (10,205 CpGs). We used the resulting components of methylation variation as independent variables in an elastic net regression using glmnet^76^.

Our hypothesis is that age acceleration is lifestyle-dependent: we therefore expect it to be bimodal for a dataset with individuals in equal parts from the zoo and the wild, which breaks model assumptions for elastic-net regression residual distribution. To overcome this limitation, we used a grid approximation approach to identify a separate best-fit slope and intercept for both groups. Namely, we divided the parameter space into an evenly spaced discrete grid, starting with possible slopes ranging from −1 to 1 in log-2 space (half to twice the ageing rate for zoo compared to wild individuals) and possible intercepts (relative age acceleration) ranging from −15 to 15 years. We also tested elastic net α mixing parameter values ranging from 0 to 1, with parameter range and step size decremented iteratively.

For each grid value, we performed a leave-one-out fit: the age of zoo individuals was first transformed linearly using the local slope and intercept, and, taking each sample out in turn, we trained an elastic net model using all principal components of methylation variance using N-fold internal cross-validation on the 63 remaining samples. We then used this model to predict epigenetic age for the 64^th^ sample. For each vector (*slope*, *intercept*, *α*), we computed the model fit as the R^2^ coefficient of a least square model where predicted age was regressed against the transformed age for each sample. We repeated this procedure iteratively at finer grid steps around the grid maximum. Finally, to evaluate the relevance of the best-fit slope and intercept, we linearly modelled predicted age as a function of true calendar age and living conditions, including individual as a random effect to account for longitudinal sampling of a subset of birds, and evaluated the magnitude and significance of the living conditions effect using estimated marginal means. For each individual, age acceleration was calculated as the difference between predicted age and true age at sampling.

This approach to define EAA was validated in two independent ways. First, we applied it to a human dataset of similar characteristics and known age acceleration parameters, including Infinium 450K BeadChip data for 64 human males of a wide range of ages, 32 of which were current smokers amongst the most age-accelerated quantile, and 32 never-smokers amongst the least age-accelerated quantile, as calculated using the independently-trained PCPhenoAge and PCGrimAge clocks^39^. We chose to focus on age-accelerated current smokers and age-decelerated non-smokers to maximise the chance that age acceleration difference between the two groups shared a physiological mechanism, in order to parallel our King penguin experimental design. The aim of this first test was to verify that our approach successfully differentiated between the two groups, and proposed age acceleration values that are consistent with well-established clocks (see Supplementary Fig. S5). Second, we repeated our approach 500 times on the King penguin dataset, randomising origin (wild/zoo) amongst individuals at each iteration, and evaluated the probability of the observed parameters against a normal distribution fit to the empirical random distribution. The aim of this second test was to test for false-positive inter-group differences (see Supplementary Fig. S4).

### Differentially methylated sites and regions

In order to identify differentially methylated sites (DMS) between wild and zoo-housed individuals, we fit per-site binomial models on observed methylated and unmethylated cytosine counts at filtered CpG sites, selected with an average depth of 10 to 25x, and an individual sample depth of 5 to 50x, with no more than 20% missing data (10,147,333 CpG sites out of 18,850,653 in the King penguin’s genome). Models were fitted using the lme4 package^77^ in R. For each model, we included as covariates the bird’s living conditions (wild or zoo), age at sampling, epigenetic age acceleration, enzymatic conversion efficiency, and genome-wide mean methylation level. Identity of the bird was included as a random factor to account for repeated sampling in some individuals. Log2(odds ratio) for living conditions was extracted from each model. We used the well-proven Metilene algorithm^78^ to identify differentially methylated regions (DMR) - a choice guided by this conservative method’s very low FDR when benchmarked against other algorithms^79^. To account for known confounding effects, however, we corrected methylation levels^80^ prior to segmentation through beta regression using glmmTMB^81^, and the same covariates as above. We retained only sites that had a minimum standard deviation of 0.1 before beta-correction, and conservatively filtered out DMRs with an absolute difference ≤ 0.1 and/or including < 10 CpG sites. Maximum distance between CpGs within a DMR was set to 500bp, corresponding to twice the median inter-CpG distance in this dataset. P-values were Benjamini-Hochberg corrected with α = 0.05. DMRs were annotated retaining only genes (including cisregulatory element) at a maximum distance of 5 kilobases from the DMR boundary. In order to evaluate the risk of false-discovery despite BH FDR correction, we conducted thorough randomisation tests (see Supplementary Fig. S9). Pathway overrepresentation was assessed using FDR-corrected binomial tests against the Reactome pathway database, using the GeneAnalytics toolkit (geneanalytics.genecards.org)^82^.

### Survival analysis

We obtained survival data for both wild and zoo King penguin populations, and calculated survival curves in both environments as described in full detail in Supplementary methods S2. Briefly, in the wild, we used capture-mark-recapture data for 1,895 known-age individuals monitored electronically since 1998^61^. Based on this data, we determined that individuals, which had not been detected for at least 2 consecutive years, had a probability of later return of ∼0.3%, and could therefore be considered dead: this information allowed us to build an empirical Bayes predictor for the posterior probability of survival. At the zoo, we collected data from the Species360 database (https://species360.org/). Data included all reliably registered King penguins held in zoos across the world since 1913 (N = 305). We retained only individuals with known hatching date, and either (i) known natural death date (N = 174) or (ii) known right-censoring (still alive) date (N = 131). Finally, we used a Cox proportional-hazard model to compare survival probability in the wild, and at the zoo (see Fig. 1C and 1D).

## Acknowledgements

This study was funded and supported by the Research Council of Finland (grants #331320 and #354649 to RC), by the French Polar Institute Paul-Emile Victor (IPEV Project 137-ANTAVIA, PI CLB), by the Centre Scientifique de Monaco (CSM) with additional support from the RTPI-NUTRESS (CSM/CNRS-UNISTRA), by the Centre National de la Recherche Scientifique (CNRS) through the Zone Atelier Antarctique et Terres Australes (ZATA), by Zoo Zurich, and by Loro Parque. This work was supported by the DFG Research Infrastructure NGS_CC (project 407495230) as part of the Next Generation Sequencing Competence Network (project 423957469). Sequencing was carried out at the Competence Centre for Genomic Analysis (Kiel). We especially acknowledge the key contribution of the Finnish IT Center for Science (CSC) for access to computational resources necessary to the realisation of this work. We also thank Prof. Jaakko Kaprio for providing data for humans used in this study. We are deeply grateful to the Zoo Zurich and Loro Parque teams for their expert support in planning and executing this experiment, as well as to all all the wintering and summering members of IPEV Project 137 and all the other colleagues and students within the P137 team, who participated in the long-term monitoring and sample collection since 2000. We also sincerely thank the IPEV logistics teams for their important and continued support in the field. This study is part of and supported by the long-term Studies in Ecology and Evolution (SEE-Life) program of the CNRS.

## Author contributions

The study was designed by RC, BM, CLB and ET. Sample collection was done by RC, CLB, LD, MEF, FG, RH, EP, GB, FANF, MH, and MO. Laboratory work was performed by RC, SZ, and SF. Analysis was designed by RC and BM, and conducted by RC, BM, and MH. Manuscript elaboration was done by all authors. All authors read and approved the final manuscript.

Supplementary Information is available for this paper.

## Supplementary material

### S0 | Study system and sampling

Peripheral blood was collected from 64 male King penguins at three locations: one in the wild (Crozet archipelago, see below) and two zoos (Zoo Zurich in Zurich, Switzerland and Loro Parque in Puerto de la Cruz, Spain). See table S1 for sample sizes and age distributions.

**(1) Crozet archipelago.** King penguins have been monitored in the wild using radio-frequency identification tags on Possession Island, Crozet Archipelago since 1998^1^. Thanks to an automated detection system, known-age individuals can be selectively recaptured at the entrance of the colony. Blood is collected periodically for a set of focal individuals, including at ∼ 10 months (when the fledglings are equipped with their subcutaneous RFID tag) and at random time points through life. Birds are captured as they leave the colony, briefly restrained for morphometric measurements, and blood is collected from the brachial vein using a 23G needle. 10 uL of whole blood is stored with 700 uL Queen’s Lysis Buffer and preserved at −20°C. In this study, we included repeated samples for 5 individuals at different time points (3 timepoints for 2 individuals, and 2 timepoints for 3 individuals - i.e. the 34 wild samples pertain to a total of 27 distinct individuals). For wild individuals, exact hatching date is not known, but synchronised breeding means the uncertainty is at most one month (nearly all hatching of viable chicks occurs between early January and early February) - a minor uncertainty compared to the considered lifespans.
**(2) Zoo Zurich.** 10 males were sampled at Zoo Zurich on May 12^th^, 2021. Blood was collected from the brachial vein and 10 uL whole blood were stored in 700 uL Queen’s Lysis Buffer. At Zoo Zurich, King penguins are held indoors, in a light- and temperature-controlled environment, with access to a swimming area. The enclosure contains only King penguins. They are fed manually with fish, supplemented with essential nutrients such as salt and vitamins, on a daily basis, and only fast spontaneously during the moulting period. No seasonal fasting is imposed on the birds.
**(3) Loro Parque.** 20 males were sampled at Loro Parque on September 5^th^, 2022. Blood was collected from the jugular vein and 10 uL whole blood were stored in 700 uL Queen’s Lysis Buffer. At Loro Parque, King penguins are held indoors, together with other penguin species, in a photoperiod- and temperature-controlled environment, enriched with artificial snow, and access to a salt-water swimming area. Feeding is organised as in Zoo Zurich, but birds are not further supplemented in salt.

**Table S1.** | Sample sizes, median age and range for the three sampling locations.

| Group | N | median age | age range |
| --- | --- | --- | --- |
| Wild - Crozet | 34 | 12 | 1 - 23 |
| Zoo - Zoo Zurich | 10 | 11 | 2.5 - 35 |
| Zoo - Loro Parque | 20 | 13 | 5 - 30 |

### S1 | DNA processing and sequencing

DNA was extracted from all blood samples using a standard spin-column protocol (Macherey-Nagel Nucleospin Blood kit), including RNAse A treatment. Samples from different origins were randomly distributed across DNA extraction batches. DNA was quantified by fluorometry. Library preparation followed NEB’s EMSeq kit protocol^2^. Libraries were sequenced on the Illumina NovaSeq 6000 at CCGA, Kiel, Germany. See Supplementary Table S3 for details.

### S2 | Survival analysis

Given the fixed-point presence-absence nature of our dataset^3^, we can strictly only estimate a right-censoring date for each individual, and not ascertain death: King penguins nearly always die at sea, so that observations of death from a land-based detection point are anecdotal. To overcome this necessary limitation, we estimated survival status of individuals in our dataset using an empirical Bayes approach. First, we restricted data to adults (at least 5 years old) who had initiated reproduction at least once in the study colony, and had attempted reproduction at least every third year since then (effectively limiting the dataset to known breeders within the study colony to avoid the confounding effect of breeding dispersal). For these, we calculated the distribution of time intervals between two detection events.

**(1) Naive classification as alive or dead.** In a first step, we aimed at classifying individuals as “likely alive” or “likely dead”, by establishing the absence pattern of an individual known to be alive. To do that, we proceeded in four steps. (*i*) We established the seasonal individual discovery curve for each year between 2001 and 2021: starting at the earliest point of seasonal return in spring (15th of October), we calculated the daily number of individuals detected that day that had not yet been detected this season (FIG S1A). (*ii*) This discovery distribution was modelled using an exponential distribution, using R package *lme4*^4^, using year and day as independent variables to explain discovery rate. We underline the fact that this model is a much better representation of the right tail of the distribution (in which we are interested) than in its left tail, which is heavily influenced by the edge effect of our procedure (all birds visiting the colony regularly around the start of the period appear in our data on the first few days of discovery, leading to artifactual steepness). Overall, the fit is satisfactory (R^2^ = 0.777, and FIG S1A). Based on that distribution, we concluded that 99.9% of the individuals that would be detected this season were expected to have already been detected by January 20th. (*iii*) For each year, taking as a reference point this “full discovery date”, we calculated the time since last detection for all individuals that were known to be alive that season (i.e. that were also detected later than this reference time point) (FIG S1B). We calculated the empirical cumulative distribution for these times since last detection, and concluded that, on average, 93.7% of individuals known to be alive on January 20th of a given year (because also detected later) had been detected within the previous 3 months, 98.8% during the previous 15 months, and 99.7% during the previous 27 months. In other words, only 0.3% birds that would later prove to be alive were ever absent from the colony for three consecutive breeding seasons (2 years and 3 months prior to the season peak on January 20th). (*iv*) Using that duration as an arbitrary threshold, we considered that a bird not detected since at least 2 years and 3 months before January 20th of the current year could be classified as “likely dead”. In this case, the bird was presumed dead during its first year of absence (since 93.7% birds return within one year to the colony) - however this is an approximation, and may lead to slightly underestimating some individuals’ lifespan.

**Fig. S1.**
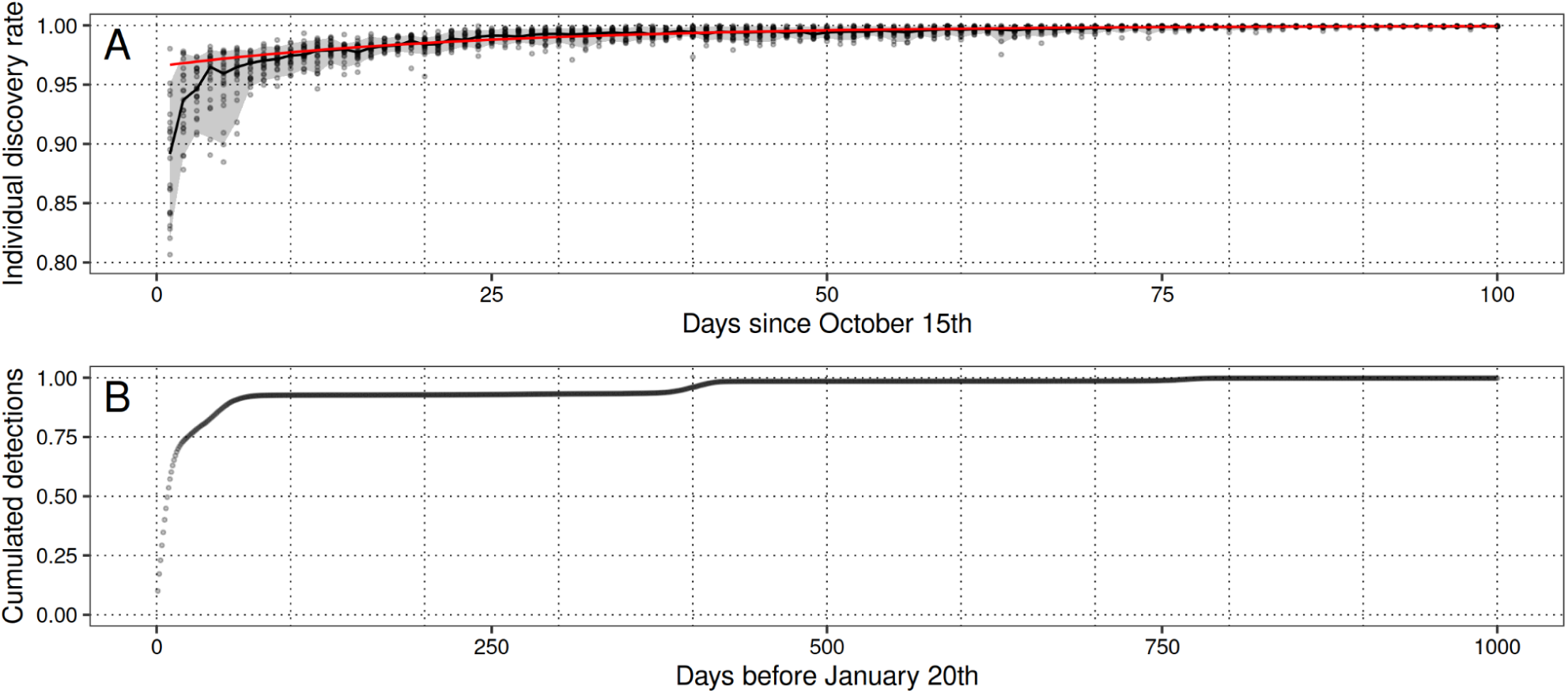
| Annual re-discovery of known individuals through time. (A) Discovery curve for the 2001-2021 period, i.e. proportion of individuals that will breed this season, and that have already been detected on a given day, computed annually starting on Oct. 15th. Shaded, median and 90% percentile interval. In red, exponential regression line. (B) Cumulative distribution function of individual resight events, for birds known to be alive, before January 20th of a given year. > 93% of living birds have normally been seen within the 3 months before January 20th (first plateau on the curve). The remaining has been detected during the previous (second plateau), or two previous breeding seasons. The plateau structure is a consequence of the absence of most adults from the colony during winter.

**(2) Survival modelling.** Based on this classification, we estimated age- and sex-specific survival probability using a Weibull regression model, using R package *survival*. Birds not seen since our classification threshold were not censored (“presumed dead”), while birds seen more recently than this threshold were right-censored (“presumed alive”). Sex and birth cohort were used as independent variables in the model.
**(3) Posterior probability of being alive.** In a second iteration, we calculated, for each individual, the posterior probability of being alive based on sex, age, and time interval since the last detection, using the canonical formulation of Bayes’ theorem (eq. 3). (*i*) *Prior probability of being alive* (eq. 1) was the Weibull-regression predicted value, given the bird’s sex and age - in other words it was the difference between age-specific survival probability at last resight, and at the reference time point (January 20th 2023). (*ii*) Probability of having been un-detected for the observed time interval given that the bird was alive (*likelihood of the “alive” model*) was derived from the observed complete distribution of inter-detection intervals, modelled using a lognormal distribution (using the “survival” package in R^5^). (*iii*) *Marginal probability of being undetected* (eq. 2) for the observed period of time was summed over both possible states, *alive* and *dead*, with the probability of the interval for a living bird given by the modelled interval distribution, and the probability of any interval for a dead bird being defined as a uniform probability density distribution between the bird’s last appearance and the reference time point.

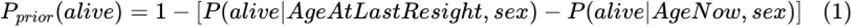

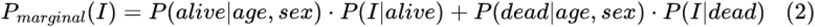

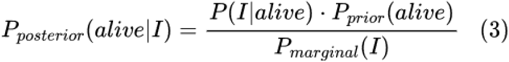
(4) Final survival modelling. We combined data from wild and zoo individuals for the final survival analysis. We used a weighted Cox PH model, using the posterior probability of being alive as evidence weight for wild birds. Each bird is included twice: once as censored, weighted by its posterior probability of being alive, and once non-censored weighted by 1 minus its posterior probability of being alive. Weight was set to 1 for zoo birds whose death had been ascertained. We included sex and rearing environment (wild or zoo) as covariates. Results and figures are presented in the main text.

### S3 | Genetic structure between groups

We tested for genetic structure between the three populations (wild birds from Crozet, and zoo birds from Zoo Zurich and Loro Parque), as epigenetic effects may be confounded by underlying genetic structure. We called single-nucleotide polymorphisms from sequencing data using the bcftools mpileup approach^6^ (v. 1.16). Since cytosines are affected by the enzymatic base-conversion protocol, we retained only AT SNPs for SNP analysis. We used vcftools^7^ (v. 0.1.16) to remove all sites with a phred-scaled quality below 40. Genotypes with a depth outside a range of 10X to 70X were excluded, and only sites in which all individuals were covered, and where the minor allele was observed at least 5 times, were retained, resulting in 486,501 AT SNPs. Five individuals were included at several age points in the analysis (see S0), only one of these samples was used for genetic analysis, reducing our dataset to 57 samples (27 from the wild and 30 from zoos).

We used PLINK2^8,9^ (v. 2.00a3) to generate a pruned dataset containing only weakly cross-correlated loci, using PLINK’s sliding-window correlation test algorithm, with a window size of 50kb by 10kb steps, and a maximum R^2^ threshold of 0.1, resulting in 147,290 unlinked SNPs. These were used to compute an identity-by-state (IBS) matrix for our samples (Fig. S2), as well as Weir and Cockerham’s pairwise F_ST_ estimator. Pairwise F_ST_ values are low between the wild and the zoo group (F_ST_ = 0.009) as well as between all three pairs of populations (Crozet / Zoo Zurich: F_ST_ = 0.022, Crozet / Loro Parque: F_ST_ = 0.013, Zoo Zurich / Loro Parque: F_ST_ = 0.034). Fixation index is very similar in Loro Parque and Crozet, with a slight excess of heterozygotes (Crozet: F = −0.045, Loro Parque: F = −0.016). There is a very slight deficit of heterozygotes in Zoo Zurich (F = 0.019). IBS is overall higher between zoo individuals than between wild individuals, which is expected given the wide difference in population sizes (as there are currently no more captures of wild King penguins for zoo rearing, all present-day zoo-kept individuals originate from the same founder group).

Further, we used fastStructure^10^ to explore possible population structure among the three sample groups, testing K values ranging from 1 to 5, with 10-fold internal cross-validation. We used fastStructure’s model complexity choice algorithm, which indicated that a single-component model (K=1) maximised the marginal likelihood, and most fully explained the structure in the data. Higher model complexities (K=2 to 5) are shown in Fig. S3: no consistent structure is inferred between populations. Overall, we consider there is no significant population structure in our data, and that genetic variation is unlikely to confound epigenetic findings.

**Fig. S2.**
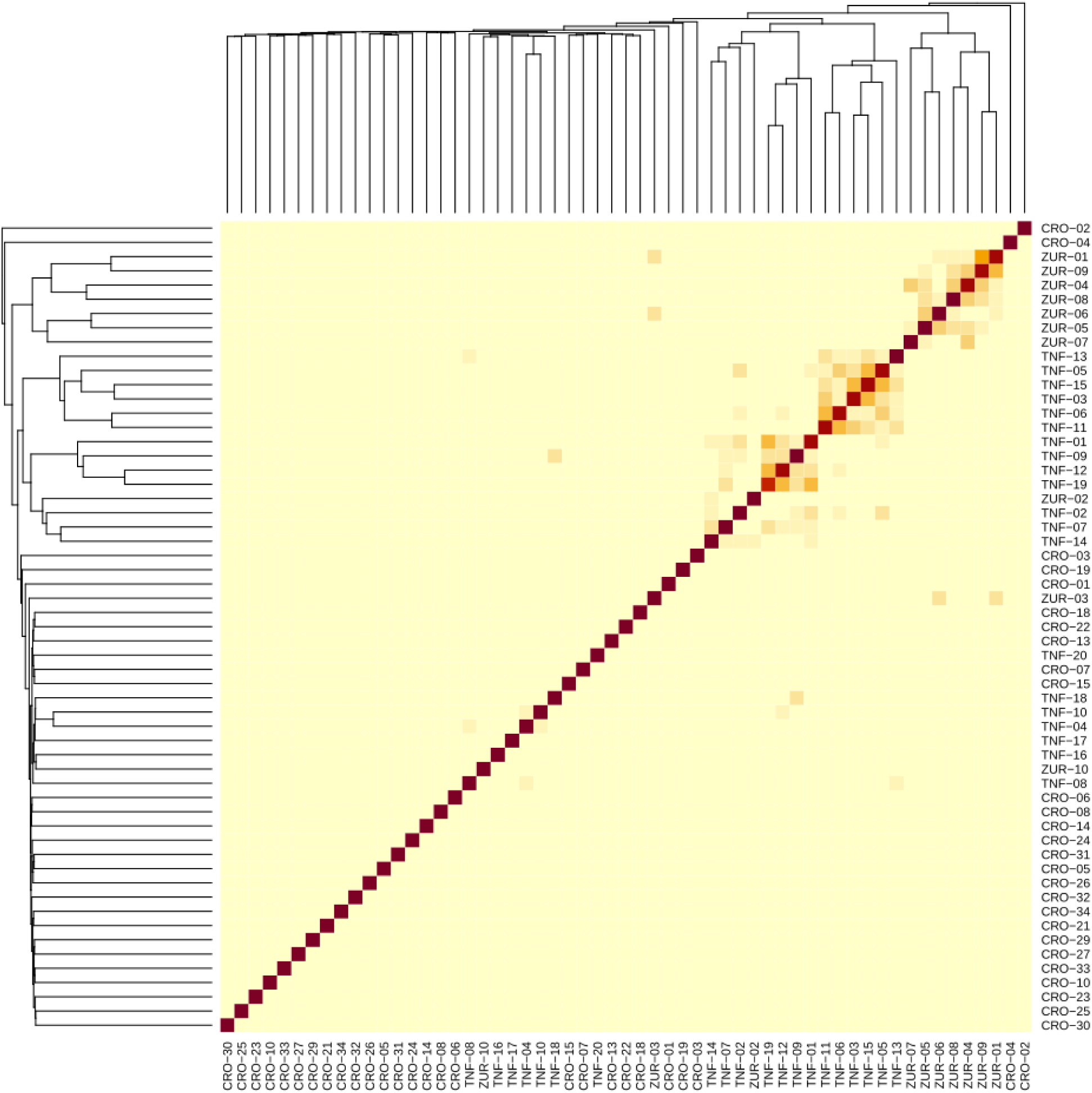
| Identity-by-state matrix, reordered by similarity. Lower genetic distances are observed between zoo individuals than between wild individuals.

**Fig. S3.**
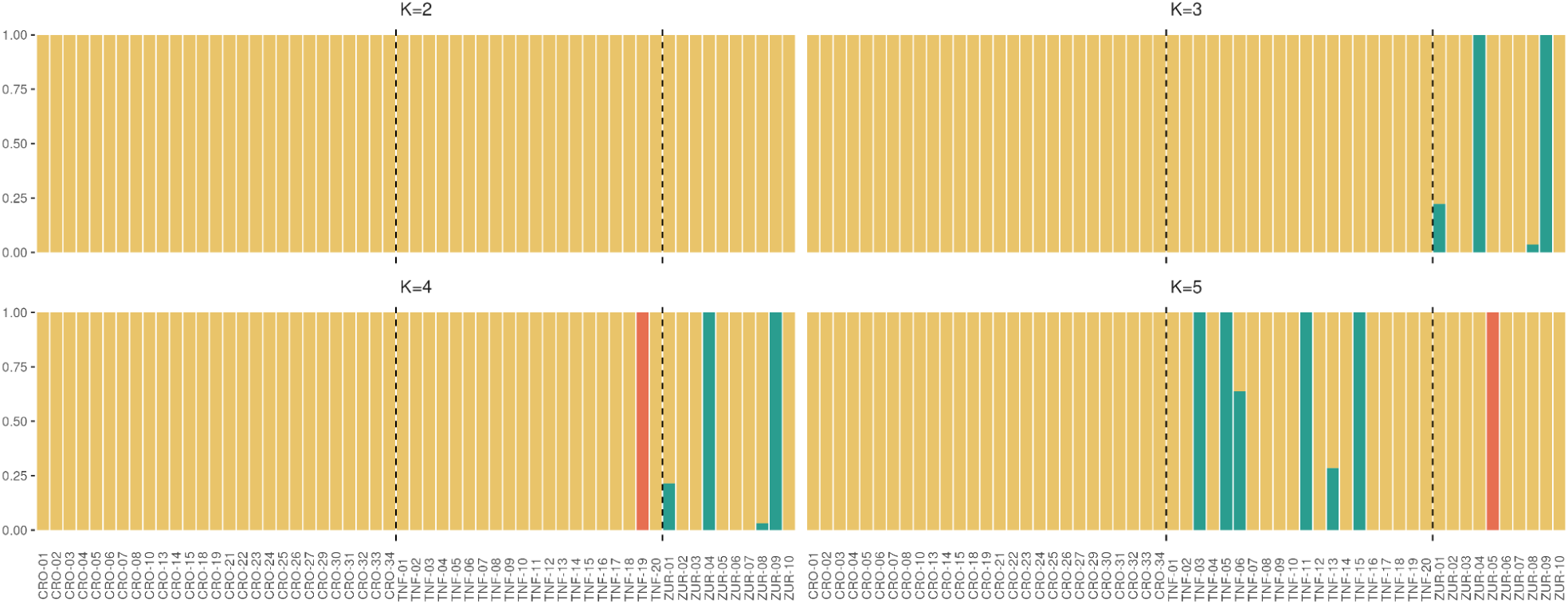
| fastStructure admixture proportions for K values from 2 to 5. Note that one or two components are always inferred to have extremely small contributions, and are not visible on the plots.

### S4 | Epigenetic age acceleration model

**Fig. S4.**
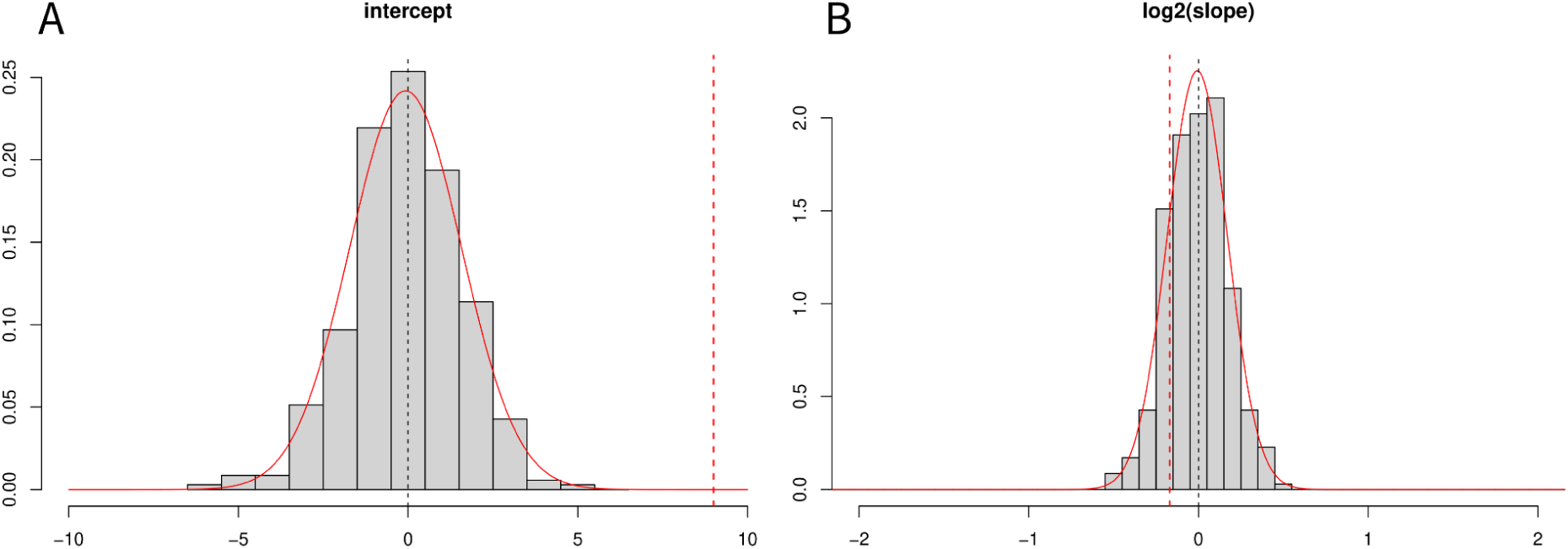
| Randomisation test for elastic net model parameters. We randomised wild/zoo rearing conditions across all samples 500 times, and repeated the entire model fitting procedure for each replicate. Distribution of intercept (A) and slope (B) parameters across replicates, together with a fitted normal distribution (red curve). The parameters inferred from the original data are displayed as vertical dashed red lines. The probability of randomly observing a value at least as different from zero (two-tailed) is approximated from the normal distribution as 3.8 x 10^−8^ ≈ 0 for the intercept, and 0.36 for the slope. This is consistent with significance estimates from linear modelling (see main text).

**Table S2.** | Modelling epigenetic age as a function of chronological age. Null model (1) excludes rearing group, while models (2) and (3) include it. Additionally, model (3) includes an interaction between the epigenetic ∼ chronological age regression slope, and the rearing group.

| (1) *epigenetic age ~ chronological age + conversion rate + genome mean* |  |  |  |  | **AIC=361.7** |
| --- | --- | --- | --- | --- | --- |
|  |  | Estimate | Standard error | t-value | p-value |
|  | *Intercept* | 169.572 | 91.960 | 1.844 | 0.0701 |
|  | *Chronological age* | 0.849 | 0.064 | 13.321 | < 2e-16 *** |
|  | *Conversion rate* | -0.683 | 0.850 | -0.803 | 0.425 |
|  | *Genome mCpG mean* | -149.686 | 42.763 | -3.500 | 0.000883 *** |
|  | *Residual standard error: 3.899 on 60 degrees of freedom* |  |  |  |  |
|  | *Multiple R-squared: 0.7879, Adjusted R-squared: 0.7773* |  |  |  |  |
|  | *F-statistic: 74.31 on 3 and 60 DF, p-value: < 2.2e-16* |  |  |  |  |
| (2) *epigenetic age ~ chronological age + rearing group + conversion rate + genome mean* |  |  |  |  | **AIC=310.1** |
|  |  | Estimate | Standard error | t-value | p-value |
|  | *Intercept (wild)* | 16.700 | 63.464 | 0.263 | 0.793 |
|  | *Chronological age* | 0.829 | 0.042 | 19.562 | < 2e-16 *** |
|  | *Group (zoo)* | 6.481 | 0.738 | 8.787 | 2.59e-12 *** |
|  | *Conversion rate* | 0.005 | 0.569 | 0.008 | 0.993 |
|  | *Genome mCpG mean* | -22.608 | 31.853 | -0.710 | 0.481 |
|  | *Residual standard error: 2.588 on 59 degrees of freedom* |  |  |  |  |
|  | *Multiple R-squared: 0.9082, Adjusted R-squared: 0.9019* |  |  |  |  |
|  | *F-statistic: 145.8 on 4 and 59 DF, p-value: < 2.2e-16* |  |  |  |  |
| (3) *epigenetic age ~ chronological age x rearing group + conversion rate + genome mean* |  |  |  |  | **AIC=308.7** |
|  |  | Estimate | Standard error | t-value | p-value |
|  | *Intercept (wild)* | 46.861 | 64.574 | 0.726 | 0.471 |
|  | *Chronological age* | 0.915 | 0.0636 | 14.385 | < 2e-16 *** |
|  | *Group (zoo)* | 8.287 | 1.245 | 6.657 | 1.1e-08 *** |
|  | *Age x Group* | -0.153 | 0.575 | -0.404 | 0.687 |
|  | *Conversion rate* | -0.232 | 31.957 | -1.073 | 0.288 |
|  | *Genome mCpG mean* | -34.285 | 0.0858 | -1.784 | 0.0797 |
|  | *Residual standard error: 2.541 on 58 degrees of freedom* |  |  |  |  |
|  | *Multiple R-squared: 0.9129, Adjusted R-squared: 0.9054* |  |  |  |  |
|  | *F-statistic: 121.6 on 5 and 58 DF, p-value: < 2.2e-16* |  |  |  |  |

#### EAA in humans as a control

We applied our approach to a human test dataset of similar characteristics (see Methods in the main text), contrasting a group of smokers and a group of non-smokers of known EAA (Supplementary Fig. S5). Our approach yielded results that were highly consistent with the independently-trained PCPhenoAge and PCGrimAge clocks^11^. Final mean difference between groups was 11.8 years according to our model (SE = 1.24, p-value < 1e^−12^), to be compared with 14.0 years according to PCPhenoAge (SE = 0.64, p-value < 1e^−16^) and 11.6 years according to PCGrimAge (SE = 0.57, p-value < 1e^−16^). Pearson’s correlation coefficient between our approach and PCPhenoAge was 0.82 (R^2^ = 0.67) and between our approach and PCGrimAge 0.75 (R^2^ = 0.57). For reference, correlation between PCPhenoAge and PCGrimAge was 0.92 (R^2^ = 0.84) - see Supplementary Fig. S5C-E. The higher accuracy of the latter two clocks is expected, given they were trained on an independent sample of 1,400 individuals, as opposed to 64 here.

**Fig. S5.**
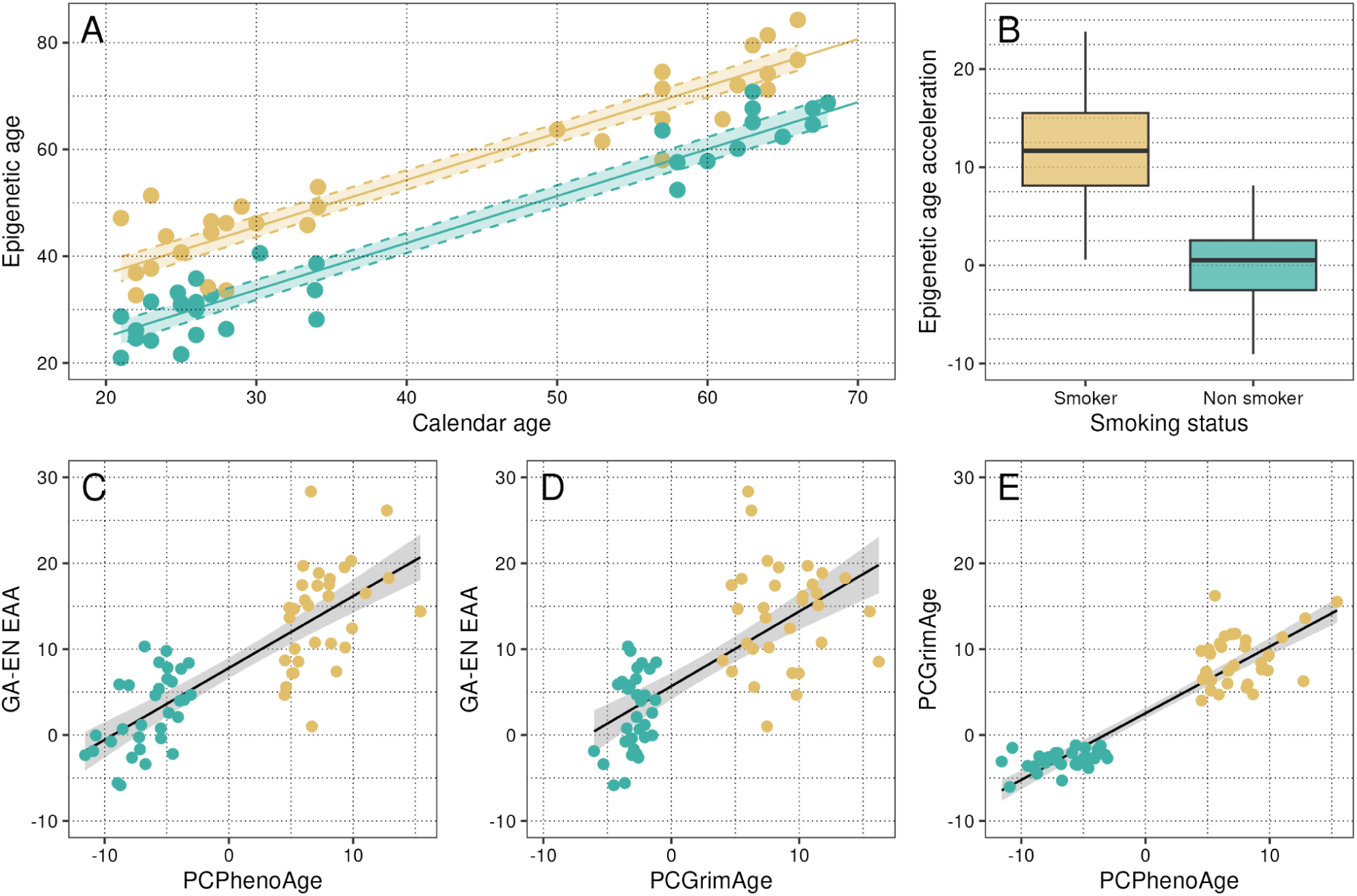
| Age acceleration in smoking and non-smoking humans. As an additional control, we applied our approach to a human dataset of similar characteristics and known age acceleration parameters, including 450K chip data for 64 Finnish human males of a wide range of ages, 32 of which were current smokers amongst the most age-accelerated quantile, and 32 never-smokers amongst the least age-accelerated quantile, as calculated using the independently-trained PCPhenoAge and PCGrimAge clocks. (A) Epigenetic age as a function of calendar age in both groups, with 95% confidence intervals. Data points are raw age predictions based on CpG methylation. (B) Distribution of age acceleration in both groups. (C), (D) and (E): comparison between our approach (Grid-approximation - Elastic net, GA-EN on the figure) and two published epigenetic clocks for the same samples. Note that PCPhenoAge and PCGrimAge were both trained on 1,400 individuals, as opposed to 64 in our GA-EN approach.

### S5 | Differentially methylated regions

We first evaluated CpG methylation across genomic features to validate the consistency of our dataset against prior knowledge. As expected, CpG methylation dropped sharply at the transcription start site (Fig. S6A), together with a peak in CpG density (Fig. S6B). Overall, CpG methylation was lower in CpG islands across features. It was lower in the first exon and the first exon of each gene, compared to subsequent introns-exons^12^. Genome-wide methylation level was slightly lower in zoo-reared penguins than in wild penguins, and this held globally across all features (average difference: 0.008, SE=0.0003, p-value <0.0001, linear model). Methodological artefacts could only arise from sample preservation conditions (as samples were randomised for library preparation and sequencing), and we consider these highly unlikely, as library preparation controls were not different between groups. Genome-wide average methylation level was, however, included as a covariate in all models, as this difference in background methylation rate would override any other differential methylation signal.

**Fig. S6.**
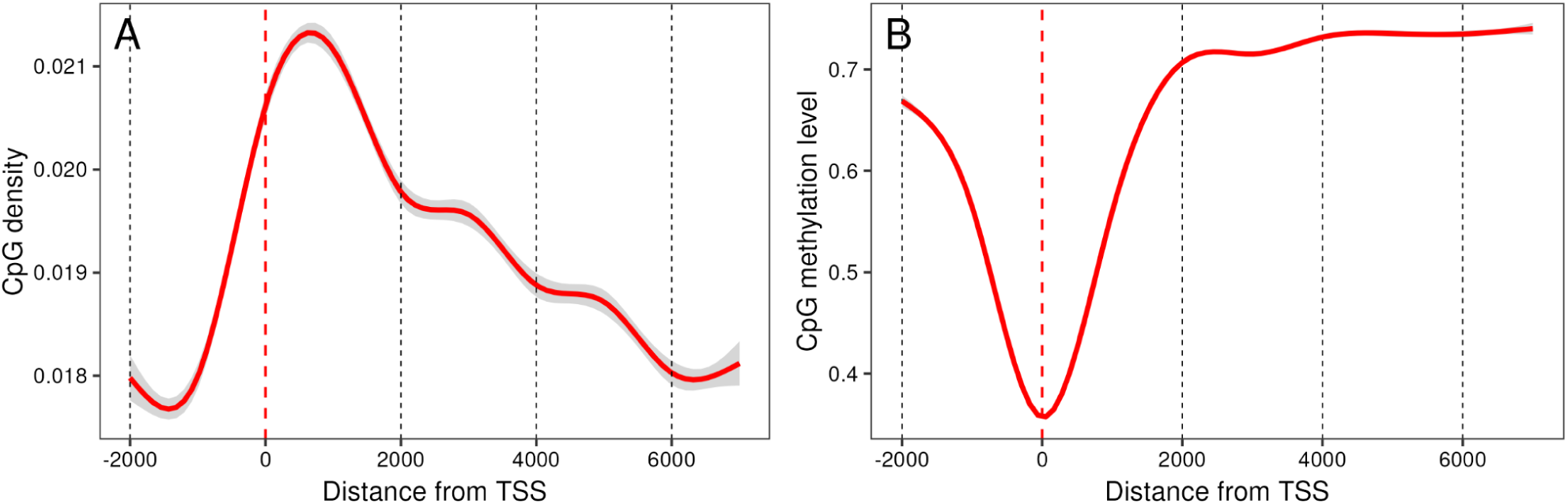
| CpG density and CpG methylation around the transcription start site. (A) Average CpG density, smoothed over 250-bp sliding windows (50bp step), around the TSS. (B) Average methylation profile around the TSS. Red line: GAM smoother, with 99% confidence interval (gray area). We considered the region starting 2kb upstream of the TSS, to 7kb downstream.

**Fig. S7.**
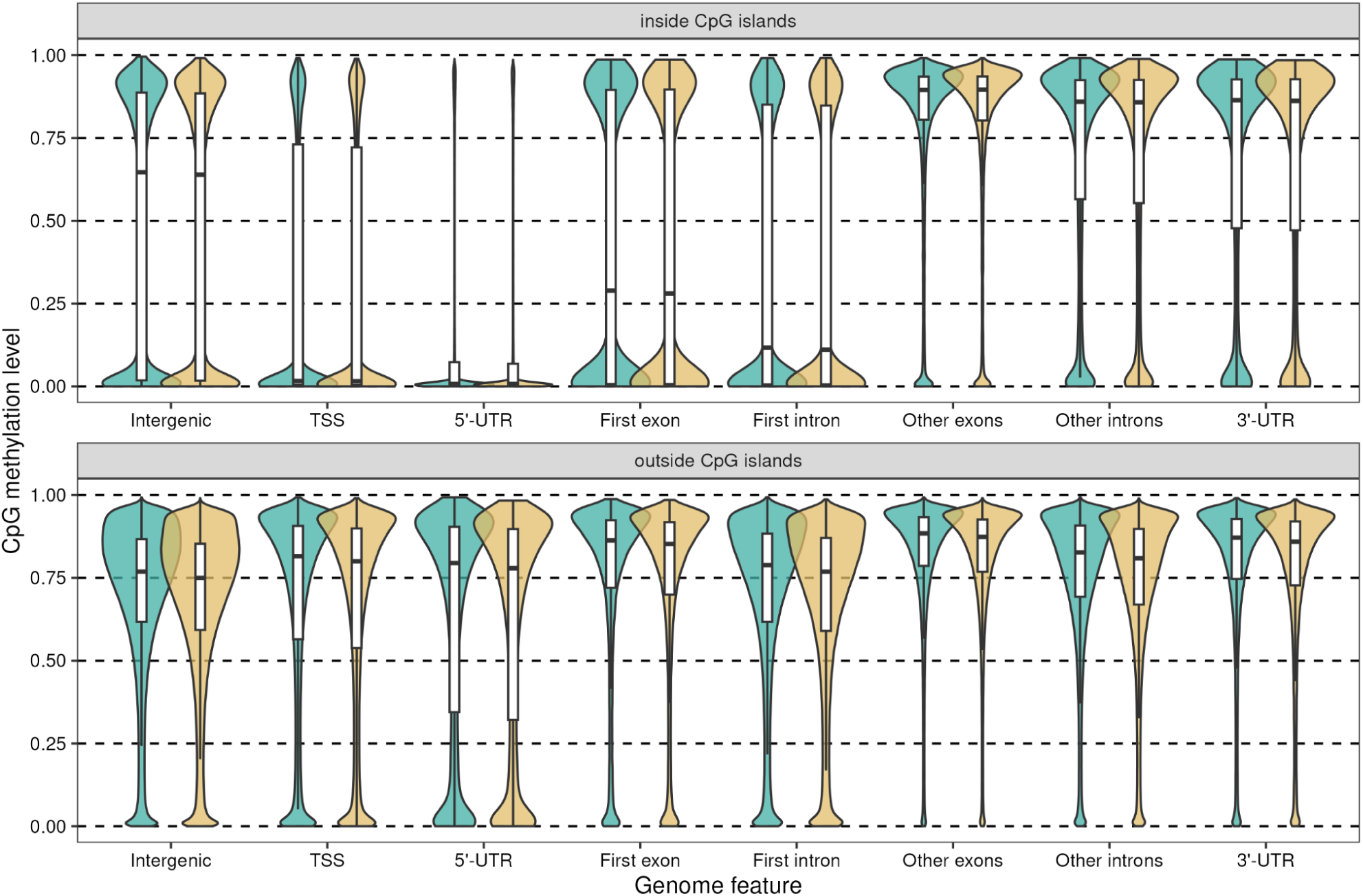
| CpG methylation across genome features, inside and outside CpG islands.

**Fig. S8.**
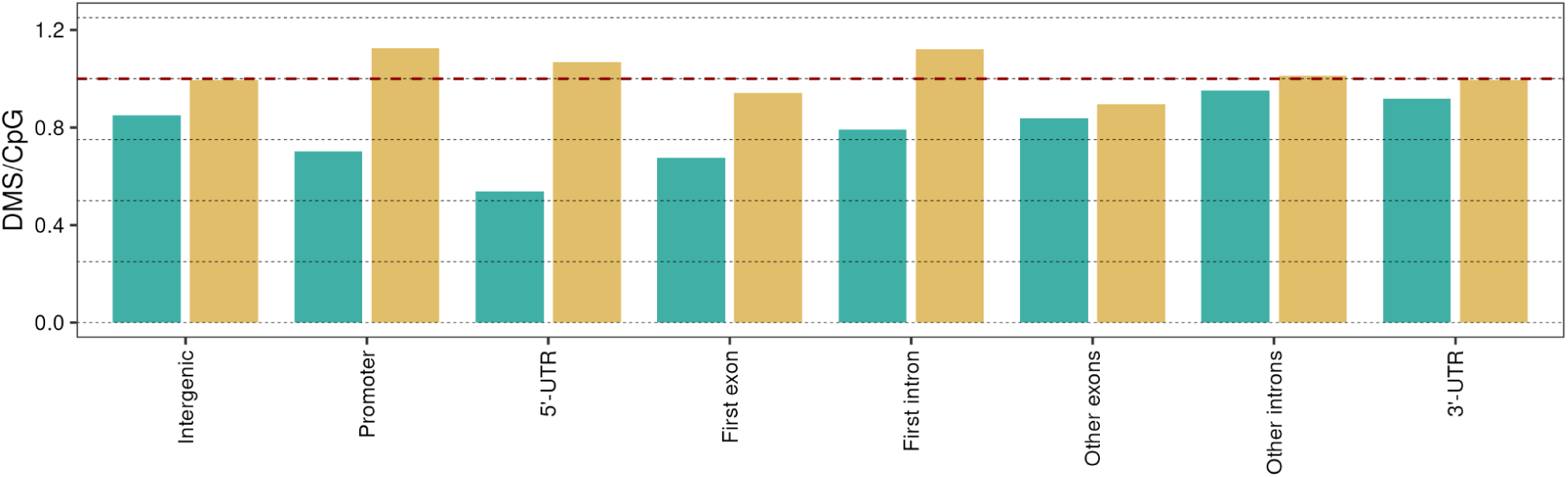
| DMS density across genome features, scaled to genome-wide DMS density. Inside CpG islands in green (left bars) and outside CpG islands (tan bars, right). DMS are defined as significant binomial tests at the 0.05 level, taking all relevant covariates into account (see main methods). Overall, we observe less DMS in CpG islands, and especially so in CpG-rich promoters (defined as above, −2000 to 7000kb from TSS), 5’-UTRs and first exons, in keeping with expectations for these mostly unmethylated areas. On the other hand, there is an excess of DMS in non-CpG-island promoters, 5’UTRs and first introns - three types of features involved in gene regulation. All pairwise tests are significant, except non-CpGi non-first exons vs non-CpGi intergenic background.

#### Metilene DMR inference

In order to evaluate the risk of false positives when using Metilene for DMR inference, we used a nested randomisation approach. First, in order to evaluate the probability of finding a false-positive DMR between random groups of birds, despite Benjamini-Hochberg FDR correction, we randomised rearing conditions across samples 500 times, and inferred DMRs based on these random assignments. We identified on average 8 DMRs in randomised data (median 7, range 0 to 40, 95% interquantile 1 to 19), to be compared with 600 DMRs inferred in the original data. Encountering a number equal or greater than 600, under a normal distribution fitted to the random distribution, would have a probability computationally indistinguishable from zero.

Second, in order to evaluate the number of genes encountered randomly within 5kbp of a dataset of 600 DMRs (as identified in our original data), we randomly shuffled these DMRs across the genome 500 times. To account for the fact that both DMRs and genes are more likely in CpG-rich areas of the genome, we shuffled DMRs only across areas of the genome that had at least as high a GC-content as our least-GC-rich DMR (i.e. a CG content ≥ 0.33). This procedure pointed to an average of 263 genes (median 263, range 233 to 295, 95% interquantile 239 to 284), to be compared with 338 in our original DMRs. Encountering a number equal or greater than 338, under a normal distribution fitted to the random distribution, would have a probability computationally ≤ 1e-10.

**Fig. S9.**
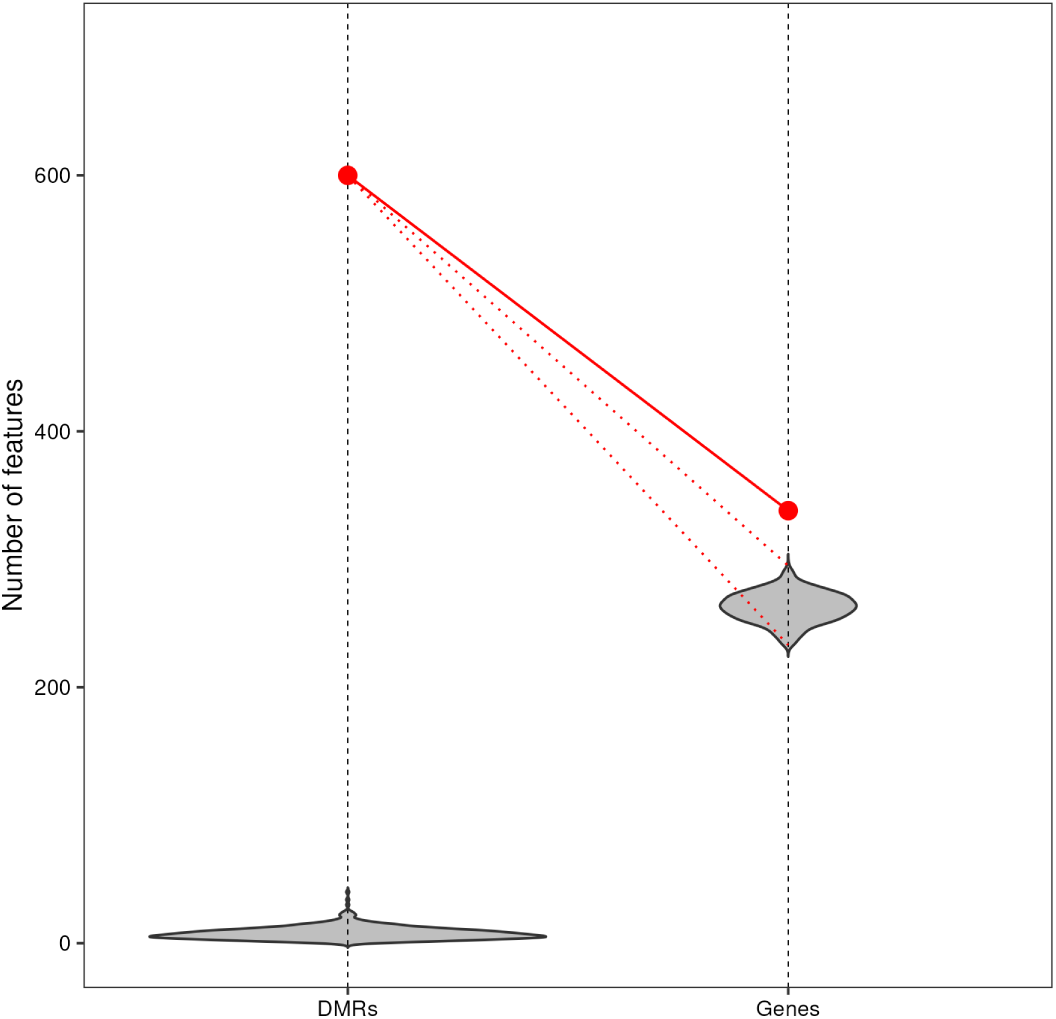
| Random DMR and differentially methylated gene discovery. Violins represent random distributions based on 500 replicates (see above). Red dots connected by a solid red line represent values inferred from the original data. The gene distribution is based on a randomisation of the 600 original DMRs (as indicated by the thin dotted lines) and not on the random-data DMRs (left violin) - in that case gene numbers are always zero.

Finally, we re-computed DMRs splitting zoo-reared birds in two groups according to their zoo of origin (Zoo Zurich, N=10 or Loro Parque, N=20) and computing DMRs for each pair of groups (each zoo vs wild, and Zoo Zurich vs Loro Parque). Numbers of DMRs were comparable in each case (737 DMRs for the wild vs. Loro Parque comparison, and 352 for the wild vs. Zoo Zurich comparison, compared to 600 for the wild vs both zoos comparison). Jaccard indices were high in both cases (0.52 for the wild vs. Loro Parque comparison, 0.21 for the wild vs. Zoo Zurich comparison - as opposed to 0.0021 [stdev. 0.0020] for 600 shuffled DMRs - see above for the randomisation protocol). When considering genes involved in each comparison, overlap between sets is high: 108 genes are involved in all three pairwise comparisons, and 190 in two pairwise comparisons (in all cases except 4, these were included in the full-set Wild vs. Zoo DMRs). Finally, 242 genes were only involved in one pairwise set of DMRs - of these possible false-positives, only 44 were uniquely found in the full Wild vs. Zoo DMRs. More noise is expected in the reduced sets, due to the reduced sample size. We consider these results to show our DMR inference is clearly out of the random expectation, and retain the full-set result for further analysis.

**Fig. S10.**
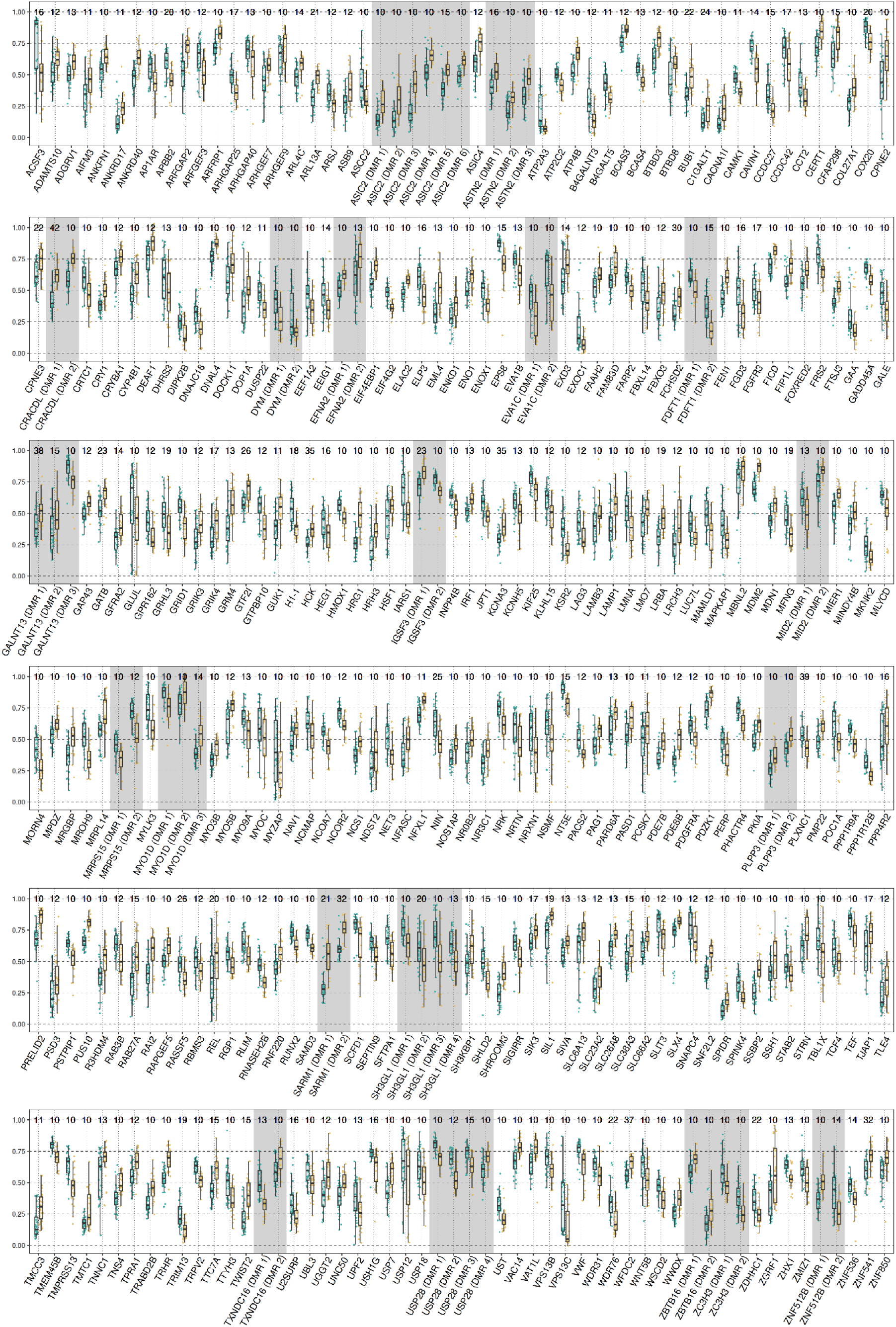
| Methylation level at all genes associated with a DMR. In green (left), wild birds, in tan, zoo birds; boxplots provide color reference. Methylation levels are beta-corrected for age, EAA and genome-wide mean CpG methylation.

## Supplementary tables S3, S4 and S5

Table S3 | Sequencing data processing metrics

Table S4 | Genes associated with DMRs.

Table S5 | Overrepresented super-paths and pathways in genes associated with DMRs

