## Supplementary tables S3-S5 for "Sedentary life accelerates epigenetic ageing in King penguins"

### Supplementary Table S3

| Name | Rearing group | Location | Age at sampling | Conversion rate | Genome mCpG mean | EAA | Raw reads | Filtered reads | Read retaining ratio | Median depth |
|---|---|---|---|---|---|---|---|---|---|---|
| A02168 | wild | Crozet | 0.86 | 97.0 | 0.65 | -3.71 | 175,545,806 | 168,005,453 | 0.96 | 20 |
| A03083 | wild | Crozet | 0.87 | 99.8 | 0.64 | -4.54 | 267,701,608 | 258,812,025 | 0.97 | 31 |
| A03374 | wild | Crozet | 0.90 | 99.8 | 0.64 | -3.16 | 291,656,096 | 277,119,813 | 0.95 | 32 |
| A03407 | wild | Crozet | 0.90 | 99.8 | 0.64 | -3.19 | 275,622,242 | 263,801,300 | 0.96 | 32 |
| KIN-BDM-2018-E-569 | wild | Crozet | 3.98 | 99.5 | 0.65 | 5.03 | 252,794,172 | 240,557,025 | 0.95 | 29 |
| KIN-BDM-2018-E-565 | wild | Crozet | 3.99 | 99.3 | 0.65 | 1.52 | 250,218,998 | 240,728,874 | 0.96 | 29 |
| KIN-BDM-2017-E-706 | wild | Crozet | 5.02 | 98.9 | 0.66 | 0.76 | 258,458,240 | 249,328,576 | 0.96 | 30 |
| KIN-BDM-2017-E-654 | wild | Crozet | 5.10 | 99.4 | 0.65 | 3.17 | 319,410,010 | 306,399,248 | 0.96 | 37 |
| A03374 | wild | Crozet | 6.04 | 99.9 | 0.65 | -0.30 | 246,801,036 | 237,441,974 | 0.96 | 28 |
| A03260 | wild | Crozet | 6.05 | 99.9 | 0.64 | -2.76 | 217,260,290 | 209,041,432 | 0.96 | 25 |
| A03407 | wild | Crozet | 6.05 | 99.8 | 0.64 | -0.66 | 206,865,944 | 198,759,692 | 0.96 | 24 |
| A03083 | wild | Crozet | 6.14 | 99.9 | 0.64 | -2.30 | 253,956,922 | 243,974,665 | 0.96 | 29 |
| KIN-BDM-2014-E-891 | wild | Crozet | 7.97 | 98.5 | 0.66 | 1.44 | 265,269,284 | 255,820,596 | 0.96 | 31 |
| KIN-BDM-2013-E-335 | wild | Crozet | 9.05 | 99.1 | 0.66 | 0.21 | 277,788,362 | 268,005,191 | 0.96 | 32 |
| E12630 | wild | Crozet | 10.01 | 99.0 | 0.66 | 1.34 | 258,707,240 | 249,314,895 | 0.96 | 30 |
| A02168 | wild | Crozet | 11.99 | 99.8 | 0.66 | 1.03 | 225,259,472 | 216,098,055 | 0.96 | 26 |
| A03083 | wild | Crozet | 11.99 | 99.9 | 0.64 | -2.52 | 258,654,808 | 248,906,127 | 0.96 | 30 |
| E10399 | wild | Crozet | 12.02 | 99.4 | 0.65 | 4.32 | 306,093,972 | 292,476,573 | 0.96 | 36 |
| A00774 | wild | Crozet | 12.20 | 99.9 | 0.64 | 1.40 | 237,172,942 | 228,822,030 | 0.96 | 27 |
| A03407 | wild | Crozet | 12.22 | 99.8 | 0.65 | -1.60 | 252,319,108 | 241,718,595 | 0.96 | 29 |
| E08115 | wild | Crozet | 14.02 | 98.9 | 0.66 | 0.79 | 175,388,374 | 168,658,705 | 0.96 | 20 |
| A03354 | wild | Crozet | 15.02 | 99.4 | 0.65 | -0.88 | 187,943,702 | 181,166,060 | 0.96 | 22 |
| A01396 | wild | Crozet | 16.03 | 99.8 | 0.65 | 0.32 | 254,900,506 | 243,062,230 | 0.95 | 29 |
| A01376 | wild | Crozet | 16.04 | 99.9 | 0.65 | 4.00 | 279,130,856 | 265,134,354 | 0.95 | 32 |
| A02580 | wild | Crozet | 17.04 | 99.3 | 0.63 | 2.36 | 244,019,280 | 234,747,619 | 0.96 | 28 |
| A01833 | wild | Crozet | 18.02 | 99.2 | 0.65 | -0.58 | 243,705,328 | 235,469,886 | 0.97 | 29 |
| A00985 | wild | Crozet | 19.02 | 99.8 | 0.65 | -1.08 | 181,227,454 | 175,790,877 | 0.97 | 21 |
| A00774 | wild | Crozet | 19.04 | 99.8 | 0.64 | -2.34 | 282,460,804 | 270,995,333 | 0.96 | 33 |
| A00388 | wild | Crozet | 19.28 | 99.8 | 0.66 | -1.53 | 242,422,950 | 232,556,553 | 0.96 | 28 |
| A00319 | wild | Crozet | 19.41 | 99.9 | 0.65 | 0.33 | 224,706,356 | 214,699,385 | 0.96 | 25 |
| A00144 | wild | Crozet | 22.12 | 99.7 | 0.65 | 1.72 | 306,030,920 | 289,497,453 | 0.95 | 35 |
| A00000 | wild | Crozet | 23.12 | 99.9 | 0.64 | 0.05 | 258,681,758 | 245,457,247 | 0.95 | 29 |
| A00017 | wild | Crozet | 23.12 | 99.5 | 0.64 | 4.93 | 234,477,890 | 223,553,639 | 0.95 | 27 |
| A00097 | wild | Crozet | 23.12 | 99.8 | 0.65 | -3.57 | 245,963,864 | 233,260,155 | 0.95 | 28 |
| LP97141 | zoo | Loro Parque | 2.51 | 99.6 | 0.65 | 6.43 | 238,569,818 | 229,294,907 | 0.96 | 27 |
| LP97139 | zoo | Loro Parque | 2.53 | 99.8 | 0.66 | 2.57 | 235,023,396 | 225,947,235 | 0.96 | 27 |
| LP97136 | zoo | Loro Parque | 3.42 | 99.2 | 0.65 | 7.00 | 279,156,730 | 266,106,892 | 0.95 | 32 |
| LP97126 | zoo | Loro Parque | 5.96 | 99.4 | 0.64 | 10.91 | 265,415,408 | 256,597,360 | 0.97 | 31 |
| LP97123 | zoo | Loro Parque | 6.57 | 99.6 | 0.65 | 7.18 | 314,165,266 | 298,299,779 | 0.95 | 36 |
| LP97120 | zoo | Loro Parque | 7.55 | 99.4 | 0.65 | 4.60 | 238,098,072 | 228,997,363 | 0.96 | 28 |
| LP97119 | zoo | Loro Parque | 8.12 | 99.7 | 0.65 | 7.23 | 261,761,848 | 251,251,262 | 0.96 | 30 |
| LP97116 | zoo | Loro Parque | 8.53 | 99.0 | 0.64 | 8.28 | 361,023,376 | 339,187,414 | 0.94 | 41 |
| LP97115 | zoo | Loro Parque | 9.48 | 99.3 | 0.63 | 10.96 | 217,395,414 | 209,375,696 | 0.96 | 25 |
| LP97114 | zoo | Loro Parque | 9.51 | 99.2 | 0.63 | 6.86 | 233,304,916 | 225,130,141 | 0.96 | 27 |
| LP97104 | zoo | Loro Parque | 12.45 | 99.6 | 0.65 | 6.15 | 239,248,174 | 230,879,525 | 0.97 | 28 |
| LP97100 | zoo | Loro Parque | 13.47 | 99.6 | 0.63 | 12.41 | 250,802,280 | 240,250,421 | 0.96 | 29 |
| LP97098 | zoo | Loro Parque | 13.51 | 99.2 | 0.64 | 3.93 | 246,817,976 | 236,917,289 | 0.96 | 28 |
| LP97091 | zoo | Loro Parque | 16.55 | 99.0 | 0.66 | 2.67 | 270,656,048 | 258,046,972 | 0.95 | 31 |
| LP97090 | zoo | Loro Parque | 16.96 | 96.6 | 0.63 | 5.81 | 231,761,348 | 221,943,054 | 0.96 | 26 |
| LP97031 | zoo | Loro Parque | 24.38 | 99.3 | 0.66 | 6.70 | 268,129,884 | 256,737,602 | 0.96 | 31 |
| LP97019 | zoo | Loro Parque | 24.39 | 99.1 | 0.64 | 7.24 | 309,771,318 | 296,302,089 | 0.96 | 36 |
| LP97010 | zoo | Loro Parque | 24.41 | 99.3 | 0.64 | 3.65 | 231,412,614 | 223,412,812 | 0.97 | 27 |
| LP97078 | zoo | Loro Parque | 28.61 | 99.5 | 0.64 | 4.48 | 246,714,716 | 236,658,624 | 0.96 | 29 |
| LP97071 | zoo | Loro Parque | 35.20 | 99.6 | 0.63 | 5.51 | 258,475,514 | 248,671,374 | 0.96 | 30 |
| Quintus | zoo | Zoo Zurich | 4.83 | 99.8 | 0.64 | 9.34 | 228,787,248 | 218,170,798 | 0.95 | 26 |
| Nils | zoo | Zoo Zurich | 7.48 | 99.9 | 0.63 | 5.90 | 271,819,610 | 255,555,678 | 0.94 | 31 |
| Noah | zoo | Zoo Zurich | 7.72 | 99.8 | 0.65 | 7.52 | 263,622,330 | 247,997,431 | 0.94 | 30 |
| Jeremy | zoo | Zoo Zurich | 11.77 | 99.8 | 0.62 | 5.57 | 286,840,628 | 270,161,303 | 0.94 | 32 |
| Ferdinand | zoo | Zoo Zurich | 12.85 | 99.8 | 0.62 | 4.72 | 331,813,222 | 308,403,938 | 0.93 | 37 |
| Engu | zoo | Zoo Zurich | 13.72 | 99.9 | 0.61 | 9.12 | 254,438,230 | 242,253,129 | 0.95 | 29 |
| Emil | zoo | Zoo Zurich | 16.76 | 99.8 | 0.62 | 6.13 | 191,547,554 | 182,378,525 | 0.95 | 21 |
| Bruno | zoo | Zoo Zurich | 16.83 | 99.8 | 0.64 | 2.39 | 271,274,620 | 257,582,919 | 0.95 | 31 |
| Seppi | zoo | Zoo Zurich | 25.85 | 99.8 | 0.62 | 4.23 | 286,687,066 | 268,465,595 | 0.94 | 32 |
| Falk | zoo | Zoo Zurich | 29.33 | 99.7 | 0.62 | 8.94 | 227,744,980 | 217,046,796 | 0.95 | 26 |

### Supplementary Table S4

| symbol | location | methylation difference | CpGs | adjusted q-value | mean methylation | | raw p-values | | assembly mapping | | |
|---|---|---|---|---|---|---|---|---|---|---|---|
| | | | | | wild | zoo | Mann-Whitney | Kolmogorov-Smirnov | scaffold | start | end |
| *ACSF3* | intron | 0.25 | 16 | 8.05E-21 | 0.76 | 0.51 | 4.59E-14 | 1.07E-25 | VULM01015654.1 | 48059 | 48552 |
| *ADAMTS10* | 5'-UTR | -0.10 | 12 | 1.47E-04 | 0.52 | 0.63 | 2.60E-12 | 1.16E-07 | VULM01011651.1 | 374706 | 375083 |
| *ADGRV1* | intron | -0.11 | 13 | 4.82E-02 | 0.47 | 0.57 | 3.22E-07 | 3.51E-04 | VULM01005296.1 | 850669 | 851801 |
| *AIFM3* | intron | -0.12 | 11 | 1.58E-03 | 0.32 | 0.45 | 1.05E-10 | 2.74E-06 | VULM01010953.1 | 121165 | 121291 |
| *ANKFN1* | intron | -0.11 | 10 | 1.86E-04 | 0.54 | 0.66 | 8.39E-13 | 1.57E-07 | VULM01006817.1 | 616056 | 616269 |
| *ANKRD17* | 5404 bp upstream of TSS | -0.10 | 11 | 7.78E-08 | 0.14 | 0.24 | 4.45E-14 | 1.19E-11 | VULM01009962.1 | 375978 | 376043 |
| *ANKRD40* | exon | -0.16 | 12 | 1.93E-15 | 0.50 | 0.66 | 4.11E-14 | 4.87E-20 | VULM01015837.1 | 214914 | 215067 |
| *AP1AR* | 1208 bp downstream of TES | 0.12 | 10 | 1.58E-02 | 0.53 | 0.41 | 7.07E-08 | 6.72E-05 | VULM01000315.1 | 56694 | 57124 |
| *APBB2* | intron | 0.13 | 20 | 1.72E-09 | 0.58 | 0.45 | 4.30E-14 | 1.68E-13 | VULM01012311.1 | 2065031 | 2065620 |
| *ARFGAP2* | 3’-UTR | -0.19 | 10 | 8.84E-14 | 0.51 | 0.70 | 3.71E-14 | 3.29E-18 | VULM01010856.1 | 1137476 | 1137640 |
| *ARFGEF3* | 723 bp upstream of TSS | 0.13 | 12 | 1.09E-03 | 0.62 | 0.49 | 3.92E-13 | 1.61E-06 | VULM01012730.1 | 391322 | 391436 |
| *ARFRP1* | intron | -0.11 | 10 | 3.88E-03 | 0.72 | 0.83 | 1.98E-11 | 9.35E-06 | VULM01004834.1 | 19059 | 19874 |
| *ARHGAP25* | 816 bp downstream of TES | 0.13 | 17 | 3.45E-11 | 0.49 | 0.36 | 4.29E-14 | 2.16E-15 | VULM01012755.1 | 113486 | 113793 |
| *ARHGAP40* | first intron | 0.11 | 13 | 5.82E-03 | 0.69 | 0.58 | 3.96E-10 | 1.62E-05 | VULM01011812.1 | 146960 | 147545 |
| *ARHGEF7* | 485 bp downstream of TES | -0.11 | 10 | 1.04E-02 | 0.44 | 0.56 | 4.89E-11 | 3.68E-05 | VULM01008382.1 | 971754 | 972001 |
| *ARHGEF9* | intron | -0.12 | 10 | 4.13E-02 | 0.61 | 0.73 | 2.83E-09 | 2.79E-04 | VULM01010275.1 | 5689986 | 5690466 |
| *ARL13A* | 136 bp downstream of TES | -0.16 | 21 | 3.67E-21 | 0.32 | 0.48 | 4.36E-14 | 4.57E-26 | VULM01008275.1 | 35441 | 35897 |
| *ARL4C* | 3'-UTR | -0.10 | 14 | 2.89E-04 | 0.47 | 0.57 | 1.36E-09 | 2.79E-07 | VULM01009119.1 | 336251 | 336914 |
| *ARSJ* | first intron | 0.10 | 12 | 2.18E-06 | 0.34 | 0.24 | 4.47E-14 | 6.46E-10 | VULM01001900.1 | 291318 | 291456 |
| *ASB9* | 5'-UTR | -0.12 | 12 | 1.77E-04 | 0.26 | 0.38 | 4.53E-14 | 1.47E-07 | VULM01015551.1 | 1346863 | 1347030 |
| *ASCC2* | 3'-UTR | 0.12 | 10 | 2.12E-02 | 0.43 | 0.31 | 4.18E-09 | 1.05E-04 | VULM01014870.1 | 3804 | 4065 |
| *ASIC2* | first intron | -0.11 | 10 | 1.40E-04 | 0.49 | 0.61 | 5.93E-11 | 1.08E-07 | VULM01008747.1 | 611365 | 611898 |
| *ASIC2* | first intron | -0.14 | 15 | 2.33E-09 | 0.40 | 0.54 | 4.62E-14 | 2.39E-13 | VULM01008747.1 | 402852 | 403587 |
| *ASIC2* | first intron | -0.11 | 10 | 4.72E-04 | 0.57 | 0.69 | 3.62E-07 | 5.31E-07 | VULM01008747.1 | 359710 | 360038 |
| *ASIC2* | first intron | -0.22 | 10 | 8.58E-24 | 0.22 | 0.43 | 4.64E-14 | 7.83E-29 | VULM01008747.1 | 351426 | 351615 |
| *ASIC2* | intron | -0.15 | 10 | 5.98E-08 | 0.18 | 0.32 | 4.07E-14 | 8.61E-12 | VULM01008747.1 | 284695 | 285055 |
| *ASIC2* | 4584 bp downstream of TES | -0.14 | 10 | 4.47E-12 | 0.16 | 0.31 | 4.43E-14 | 2.33E-16 | VULM01008747.1 | 275855 | 276179 |
| *ASIC4* | 5759 bp downstream of TES | -0.16 | 12 | 4.34E-06 | 0.61 | 0.76 | 4.12E-14 | 1.50E-09 | VULM01012923.1 | 77448 | 77666 |
| *ASTN2* | intron | -0.12 | 16 | 1.37E-06 | 0.40 | 0.52 | 4.13E-14 | 3.78E-10 | VULM01013715.1 | 1326386 | 1326614 |
| *ASTN2* | intron | -0.11 | 10 | 6.35E-08 | 0.20 | 0.31 | 4.62E-14 | 9.28E-12 | VULM01013715.1 | 1411461 | 1411729 |
| *ASTN2* | intron | -0.14 | 10 | 3.55E-07 | 0.32 | 0.46 | 4.74E-14 | 7.49E-11 | VULM01013715.1 | 1418600 | 1418848 |
| *ATP2A3* | intron | 0.12 | 10 | 9.84E-06 | 0.24 | 0.12 | 6.99E-14 | 4.09E-09 | VULM01003902.1 | 1100746 | 1101222 |
| *ATP2C2* | intron | 0.10 | 12 | 1.02E-03 | 0.53 | 0.43 | 1.36E-08 | 1.48E-06 | VULM01009081.1 | 724475 | 724599 |
| *ATP4B* | intron | -0.13 | 10 | 5.04E-03 | 0.56 | 0.69 | 1.68E-09 | 1.33E-05 | VULM01003239.1 | 2580608 | 2580904 |
| *B4GALNT3* | 687 bp upstream of TSS | 0.15 | 12 | 2.63E-14 | 0.33 | 0.18 | 4.07E-14 | 8.67E-19 | VULM01001894.1 | 4939569 | 4940007 |
| *B4GALT5* | 3868 bp downstream of TES | 0.11 | 11 | 7.99E-06 | 0.41 | 0.30 | 5.37E-14 | 3.18E-09 | VULM01009715.1 | 1336107 | 1336266 |
| *BCAS3* | intron | -0.10 | 12 | 1.24E-08 | 0.76 | 0.87 | 4.13E-14 | 1.50E-12 | VULM01000654.1 | 2628297 | 2628433 |
| *BCAS4* | 6378 bp upstream of TSS | 0.11 | 13 | 1.35E-04 | 0.59 | 0.47 | 3.84E-13 | 1.03E-07 | VULM01006415.1 | 389150 | 389356 |
| *BTBD3* | exon | -0.16 | 12 | 2.34E-12 | 0.63 | 0.79 | 4.03E-14 | 1.13E-16 | VULM01007866.1 | 2511362 | 2511665 |
| *BTBD8* | 5'-UTR | -0.11 | 10 | 1.41E-03 | 0.46 | 0.57 | 6.44E-09 | 2.31E-06 | VULM01011373.1 | 524913 | 525337 |
| *BUB1* | 5421 bp downstream of TES | -0.16 | 22 | 5.57E-10 | 0.34 | 0.50 | 4.42E-14 | 4.44E-14 | VULM01009959.1 | 263187 | 263746 |
| *C1GALT1* | 5'-UTR | -0.10 | 24 | 1.77E-14 | 0.14 | 0.24 | 4.27E-14 | 5.36E-19 | VULM01002945.1 | 63422 | 64096 |
| *CACNA1I* | intron | -0.10 | 10 | 2.83E-03 | 0.17 | 0.28 | 1.22E-09 | 6.10E-06 | VULM01000896.1 | 431041 | 431431 |
| *CAMK1* | first intron | 0.11 | 11 | 1.27E-03 | 0.46 | 0.35 | 2.27E-10 | 1.98E-06 | VULM01006483.1 | 2152086 | 2152481 |
| *CAVIN1* | first intron | 0.19 | 14 | 1.74E-17 | 0.72 | 0.53 | 4.13E-14 | 3.30E-22 | VULM01009776.1 | 43757 | 43976 |
| *CCDC27* | first intron | 0.12 | 15 | 6.92E-05 | 0.35 | 0.23 | 4.15E-14 | 4.39E-08 | VULM01010605.1 | 1653188 | 1653486 |
| *CCDC42* | 5'-UTR to first intron | 0.13 | 17 | 3.73E-08 | 0.69 | 0.57 | 4.59E-14 | 5.10E-12 | VULM01011802.1 | 2435462 | 2435655 |
| *CCT2* | 508 bp downstream of TES | 0.12 | 13 | 2.54E-04 | 0.44 | 0.32 | 2.16E-13 | 2.35E-07 | VULM01012635.1 | 518001 | 518756 |
| *CERT1* | 1064 bp downstream of TES | -0.11 | 10 | 2.04E-02 | 0.73 | 0.84 | 1.47E-10 | 9.84E-05 | VULM01006363.1 | 802194 | 802609 |
| *CFAP298* | 3'-UTR | -0.15 | 15 | 1.82E-17 | 0.64 | 0.79 | 4.93E-14 | 3.58E-22 | VULM01014838.1 | 1083735 | 1083993 |
| *COL27A1* | first intron | -0.11 | 10 | 1.38E-03 | 0.33 | 0.45 | 4.98E-09 | 2.26E-06 | VULM01000193.1 | 205747 | 206126 |
| *COX20* | 3'-UTR | 0.12 | 20 | 3.07E-29 | 0.88 | 0.76 | 5.46E-14 | 1.44E-34 | VULM01003756.1 | 2775954 | 2776265 |
| *CPNE2* | 3537 bp downstream of TES | -0.19 | 10 | 4.20E-07 | 0.45 | 0.64 | 4.22E-14 | 9.40E-11 | VULM01010374.1 | 503963 | 504049 |
| *CPNE3* | last intron to 3'-UTR | -0.11 | 22 | 2.08E-04 | 0.62 | 0.73 | 4.45E-14 | 1.82E-07 | VULM01007254.1 | 1211186 | 1213160 |
| *CRACDL* | 4746 bp upstream of TSS | -0.15 | 10 | 5.43E-06 | 0.61 | 0.77 | 4.21E-14 | 1.97E-09 | VULM01007759.1 | 6079676 | 6079976 |
| *CRACDL* | 2743 bp upstream of TSS | -0.19 | 42 | 2.30E-35 | 0.43 | 0.62 | 0.00E+00 | 4.85E-41 | VULM01007759.1 | 6077673 | 6079039 |
| *CRTC1* | 2052 bp upstream of TSS | 0.13 | 10 | 6.66E-06 | 0.62 | 0.49 | 4.26E-14 | 2.56E-09 | VULM01015498.1 | 49781 | 49989 |
| *CRY1* | intron | -0.11 | 10 | 1.27E-02 | 0.43 | 0.55 | 1.11E-06 | 4.92E-05 | VULM01006140.1 | 3116274 | 3117122 |
| *CRYBA1* | 1075 bp downstream of TES | -0.11 | 12 | 6.36E-05 | 0.66 | 0.77 | 7.76E-14 | 3.94E-08 | VULM01000933.1 | 10365 | 10533 |
| *CYP4B1* | 4838 bp downstream of TES | -0.13 | 10 | 1.46E-04 | 0.49 | 0.62 | 8.12E-13 | 1.14E-07 | VULM01011359.1 | 37676 | 38066 |
| *DEAF1* | 1568 bp downstream of TES | -0.12 | 12 | 2.06E-11 | 0.75 | 0.87 | 4.66E-14 | 1.25E-15 | VULM01013076.1 | 148907 | 149026 |
| *DHRS3* | 4462 bp downstream of TES | 0.11 | 13 | 4.72E-02 | 0.58 | 0.46 | 1.58E-10 | 3.40E-04 | VULM01014951.1 | 778229 | 778333 |
| *DIPK2B* | first intron | 0.11 | 10 | 2.12E-04 | 0.26 | 0.15 | 6.51E-11 | 1.86E-07 | VULM01015068.1 | 1034677 | 1034950 |
| *DNAJC18* | intron | 0.11 | 10 | 9.09E-08 | 0.33 | 0.22 | 4.10E-14 | 1.40E-11 | VULM01010858.1 | 3733 | 3875 |
| *DNAL4* | 1020 bp downstream of TES | -0.10 | 10 | 3.90E-08 | 0.78 | 0.88 | 4.72E-14 | 5.40E-12 | VULM01015280.1 | 27512 | 27776 |
| *DOCK11* | intron | -0.13 | 10 | 4.37E-07 | 0.56 | 0.69 | 4.66E-14 | 9.85E-11 | VULM01004504.1 | 633927 | 634088 |
| *DOP1A* | intron-exon boundary | -0.13 | 12 | 1.11E-07 | 0.37 | 0.49 | 4.67E-14 | 1.80E-11 | VULM01003851.1 | 84832 | 85287 |
| *DUSP22* | first intron | 0.15 | 11 | 2.28E-09 | 0.51 | 0.35 | 4.31E-14 | 2.32E-13 | VULM01012134.1 | 976 | 1043 |
| *DYM* | intron | 0.14 | 10 | 1.66E-04 | 0.42 | 0.28 | 9.63E-14 | 1.37E-07 | VULM01007136.1 | 2324364 | 2324759 |
| *DYM* | intron | 0.13 | 10 | 4.58E-05 | 0.29 | 0.16 | 4.25E-14 | 2.59E-08 | VULM01007136.1 | 2370126 | 2370293 |
| *EEF1A2* | 394 bp upstream of TSS | 0.12 | 10 | 1.20E-03 | 0.47 | 0.35 | 2.71E-11 | 1.84E-06 | VULM01002311.1 | 1534586 | 1534762 |
| *EEIG1* | 3'-UTR | 0.12 | 14 | 3.12E-06 | 0.47 | 0.35 | 4.27E-14 | 9.75E-10 | VULM01000193.1 | 439928 | 440038 |
| *EFNA2* | first intron | -0.17 | 13 | 8.72E-13 | 0.59 | 0.76 | 4.53E-14 | 3.98E-17 | VULM01008410.1 | 71020 | 71235 |
| *EFNA2* | intron | -0.12 | 10 | 9.21E-03 | 0.52 | 0.64 | 4.98E-07 | 3.12E-05 | VULM01008410.1 | 47381 | 47741 |
| *EIF4EBP1* | 5'-UTR | -0.15 | 10 | 4.28E-12 | 0.55 | 0.70 | 4.50E-14 | 2.17E-16 | VULM01014478.1 | 82864 | 83076 |
| *EIF4G2* | 810 bp upstream of TSS | 0.12 | 10 | 1.16E-06 | 0.47 | 0.35 | 1.03E-10 | 3.07E-10 | VULM01009891.1 | 542203 | 542543 |
| *ELAC2* | intron-exon boundary | -0.11 | 10 | 8.72E-03 | 0.50 | 0.61 | 3.38E-06 | 2.88E-05 | VULM01012313.1 | 1546142 | 1546807 |
| *ELP3* | intron | 0.14 | 16 | 1.47E-05 | 0.59 | 0.45 | 4.56E-14 | 6.54E-09 | VULM01001811.1 | 1651231 | 1651569 |
| *EML4* | 6258 bp upstream of TSS | -0.20 | 13 | 1.05E-07 | 0.32 | 0.52 | 4.43E-14 | 1.68E-11 | VULM01008661.1 | 2086304 | 2086546 |
| *ENKD1* | 5’-UTR | -0.11 | 10 | 1.61E-02 | 0.26 | 0.37 | 9.92E-08 | 6.91E-05 | VULM01010374.1 | 654685 | 654906 |
| *ENO1* | 3'-UTR | -0.10 | 10 | 1.25E-02 | 0.48 | 0.59 | 4.74E-08 | 4.81E-05 | VULM01008382.1 | 2143648 | 2143844 |
| *ENOX1* | intron | 0.10 | 10 | 3.27E-03 | 0.50 | 0.40 | 7.61E-08 | 7.41E-06 | VULM01005679.1 | 3734539 | 3734824 |
| *EPS8* | 3’-UTR | 0.19 | 15 | 1.17E-31 | 0.88 | 0.69 | 5.26E-14 | 3.30E-37 | VULM01001894.1 | 2103949 | 2104323 |
| *EVA1B* | 2337 bp upstream of TSS | 0.10 | 13 | 3.19E-04 | 0.72 | 0.62 | 4.54E-14 | 3.19E-07 | VULM01012774.1 | 66561 | 66689 |
| *EVA1C* | 3’-UTR | 0.14 | 10 | 3.92E-02 | 0.40 | 0.26 | 4.16E-08 | 2.58E-04 | VULM01014838.1 | 1058080 | 1058567 |
| *EVA1C* | 3’-UTR | 0.19 | 10 | 7.14E-06 | 0.62 | 0.42 | 5.41E-14 | 2.77E-09 | VULM01014838.1 | 1058597 | 1059350 |
| *EXD3* | intron | -0.12 | 14 | 2.36E-04 | 0.61 | 0.73 | 4.29E-13 | 2.14E-07 | VULM01015592.1 | 270669 | 270948 |
| *EXOC1* | 477 bp upstream of TSS | 0.11 | 12 | 1.93E-04 | 0.25 | 0.14 | 1.27E-08 | 1.64E-07 | VULM01004407.1 | 1623950 | 1624179 |
| *FAAH2* | 5459 bp upstream of TSS | -0.12 | 10 | 3.02E-04 | 0.50 | 0.63 | 4.91E-14 | 2.96E-07 | VULM01015134.1 | 63432 | 63550 |
| *FAM83D* | intron-exon boundary | -0.13 | 10 | 2.53E-04 | 0.57 | 0.70 | 1.04E-10 | 2.34E-07 | VULM01011812.1 | 9277 | 10622 |
| *FARP2* | intron | 0.12 | 10 | 7.73E-04 | 0.62 | 0.50 | 5.92E-09 | 1.03E-06 | VULM01002609.1 | 746455 | 747257 |
| *FBXL14* | 4356 bp downstream of TES | 0.12 | 10 | 5.90E-04 | 0.57 | 0.45 | 1.68E-10 | 7.14E-07 | VULM01001894.1 | 4330197 | 4330351 |
| *FBXO3* | 1414 bp downstream of TES | -0.13 | 12 | 1.33E-05 | 0.35 | 0.48 | 4.22E-14 | 5.85E-09 | VULM01015328.1 | 6019931 | 6020134 |
| *FCHSD2* | 3'-UTR | -0.12 | 30 | 6.61E-12 | 0.33 | 0.45 | 4.05E-14 | 3.62E-16 | VULM01011004.1 | 550630 | 551214 |
| *FDFT1* | 4387 bp upstream of TSS | 0.17 | 15 | 5.13E-15 | 0.36 | 0.19 | 4.91E-14 | 1.37E-19 | VULM01013669.1 | 1068938 | 1069271 |
| *FDFT1* | 3717 bp upstream of TSS | 0.12 | 10 | 2.46E-02 | 0.62 | 0.49 | 1.37E-09 | 1.30E-04 | VULM01013669.1 | 1068268 | 1068639 |
| *FEN1* | 236 bp downstream of TES | -0.17 | 10 | 4.19E-04 | 0.44 | 0.61 | 2.09E-12 | 4.55E-07 | VULM01002076.1 | 61776 | 62014 |
| *FGD3* | intron-exon boundary | 0.17 | 16 | 7.41E-13 | 0.49 | 0.32 | 4.44E-14 | 3.33E-17 | VULM01010079.1 | 277837 | 278329 |
| *FGFR3* | 3490 bp upstream of TSS | 0.10 | 17 | 6.83E-05 | 0.50 | 0.40 | 4.42E-14 | 4.32E-08 | VULM01013333.1 | 3760738 | 3760905 |
| *FICD* | first intron | -0.11 | 10 | 6.97E-06 | 0.71 | 0.81 | 6.11E-14 | 2.69E-09 | VULM01005923.1 | 870170 | 870390 |
| *FIP1L1* | intron | -0.12 | 10 | 1.03E-02 | 0.55 | 0.68 | 4.81E-07 | 3.68E-05 | VULM01010275.1 | 5189900 | 5190314 |
| *FOXRED2* | 3'-UTR | -0.12 | 10 | 8.28E-03 | 0.55 | 0.67 | 4.23E-11 | 2.67E-05 | VULM01006140.1 | 1049547 | 1049688 |
| *FRS2* | 3733 bp downstream of TES | 0.13 | 10 | 1.95E-14 | 0.79 | 0.66 | 3.67E-14 | 6.15E-19 | VULM01012635.1 | 568691 | 568800 |
| *FTSJ3* | intron-exon boundary | -0.12 | 10 | 1.01E-02 | 0.43 | 0.55 | 1.59E-08 | 3.54E-05 | VULM01008747.1 | 179829 | 180067 |
| *GAA* | intron | 0.10 | 10 | 6.66E-03 | 0.31 | 0.21 | 1.19E-08 | 1.95E-05 | VULM01015262.1 | 5453 | 5684 |
| *GADD45A* | 1103 bp downstream of TES | 0.10 | 10 | 4.81E-02 | 0.66 | 0.56 | 7.05E-08 | 3.50E-04 | VULM01015833.1 | 895744 | 896086 |
| *GALE* | intron-exon boundary | 0.13 | 10 | 6.75E-04 | 0.47 | 0.35 | 2.70E-10 | 8.58E-07 | VULM01007308.1 | 2707 | 2832 |
| *GALNT13* | 928 bp upstream of TSS | -0.11 | 38 | 2.57E-10 | 0.40 | 0.51 | 4.23E-14 | 1.91E-14 | VULM01013455.1 | 639840 | 640933 |
| *GALNT13* | intron | 0.12 | 10 | 1.26E-14 | 0.86 | 0.74 | 4.89E-14 | 3.63E-19 | VULM01013455.1 | 768025 | 768132 |
| *GALNT13* | 708 bp downstream of TES | -0.13 | 15 | 2.78E-04 | 0.37 | 0.50 | 5.05E-12 | 2.66E-07 | VULM01013455.1 | 642730 | 643551 |
| *GAP43* | 749 bp downstream of TES | -0.10 | 12 | 9.29E-04 | 0.48 | 0.58 | 2.73E-10 | 1.30E-06 | VULM01006677.1 | 3824396 | 3824531 |
| *GATB* | intron | -0.13 | 23 | 6.46E-06 | 0.52 | 0.65 | 4.06E-14 | 2.46E-09 | VULM01004360.1 | 816636 | 817380 |
| *GFRA2* | 4350 bp downstream of TES | -0.11 | 14 | 3.39E-07 | 0.32 | 0.43 | 4.69E-14 | 6.99E-11 | VULM01015196.1 | 121290 | 121634 |
| *GLUL* | 2685 bp upstream of TSS | 0.10 | 10 | 2.82E-02 | 0.53 | 0.43 | 9.44E-05 | 1.59E-04 | VULM01003513.1 | 2951580 | 2951882 |
| *GPR162* | intron | 0.12 | 12 | 7.87E-04 | 0.42 | 0.30 | 4.09E-12 | 1.05E-06 | VULM01009205.1 | 427564 | 428468 |
| *GRHL3* | 132 bp downstream of TES | 0.12 | 19 | 6.93E-15 | 0.51 | 0.39 | 4.31E-14 | 1.95E-19 | VULM01007190.1 | 165511 | 165787 |
| *GRID1* | first intron | 0.14 | 10 | 1.36E-07 | 0.53 | 0.39 | 4.07E-14 | 2.37E-11 | VULM01013999.1 | 1135486 | 1135585 |
| *GRIK3* | first intron | -0.14 | 12 | 6.13E-10 | 0.30 | 0.43 | 4.23E-14 | 5.13E-14 | VULM01012774.1 | 339373 | 339496 |
| *GRIK4* | intron | -0.11 | 17 | 4.65E-04 | 0.30 | 0.42 | 4.43E-14 | 5.20E-07 | VULM01014095.1 | 101463 | 101987 |
| *GRM4* | intron | -0.20 | 13 | 3.85E-07 | 0.40 | 0.59 | 4.41E-14 | 8.34E-11 | VULM01014263.1 | 31933 | 32150 |
| *GTF2I* | 6198 bp upstream of TSS | -0.11 | 26 | 2.99E-07 | 0.57 | 0.68 | 4.63E-14 | 6.02E-11 | VULM01003902.1 | 1885954 | 1887241 |
| *GTPBP10* | 634 bp upstream of TSS | 0.16 | 12 | 9.76E-07 | 0.55 | 0.39 | 4.53E-14 | 2.50E-10 | VULM01006696.1 | 3172503 | 3172736 |
| *GUK1* | 1142 bp downstream of TES | -0.15 | 11 | 2.81E-05 | 0.42 | 0.57 | 4.29E-14 | 1.43E-08 | VULM01011723.1 | 152490 | 152660 |
| *H1-1* | 1286 bp upstream of TSS | 0.16 | 18 | 1.95E-14 | 0.54 | 0.38 | 4.64E-14 | 6.17E-19 | VULM01001160.1 | 49668 | 50184 |
| *HCK* | 3'-UTR | -0.12 | 35 | 5.67E-15 | 0.26 | 0.39 | 3.86E-14 | 1.55E-19 | VULM01014051.1 | 494 | 1137 |

| | | | | | | | | | | | |
|---|---|---|---|---|---|---|---|---|---|---|---|
| *HEG1* | intron-exon boundary | 0.10 | 16 | 7.91E-03 | 0.44 | 0.34 | 1.33E-07 | 2.49E-05 | VULM01012013.1 | 753922 | 754423 |
| *HMOX1* | 2865 bp upstream of TSS | 0.10 | 10 | 2.28E-02 | 0.51 | 0.41 | 2.55E-05 | 1.17E-04 | VULM01006140.1 | 1505396 | 1505872 |
| *HRG1* | 5'-UTR | -0.21 | 10 | 6.32E-09 | 0.29 | 0.50 | 4.46E-14 | 7.02E-13 | VULM01000284.1 | 59837 | 60161 |
| *HRH3* | first intron | -0.15 | 10 | 5.57E-10 | 0.22 | 0.37 | 4.74E-14 | 4.46E-14 | VULM01003799.1 | 3607034 | 3607137 |
| *HSF1* | first intron | -0.12 | 10 | 2.61E-04 | 0.45 | 0.56 | 1.09E-08 | 2.45E-07 | VULM01014010.1 | 148796 | 149112 |
| *IARS1* | 745 bp downstream of TES | 0.19 | 10 | 6.04E-06 | 0.66 | 0.47 | 1.30E-12 | 2.24E-09 | VULM01013496.1 | 159655 | 160396 |
| *IGSF3* | intron | 0.11 | 10 | 9.75E-08 | 0.78 | 0.67 | 4.19E-14 | 1.55E-11 | VULM01014500.1 | 175331 | 176211 |
| *IGSF3* | intron-exon boundary | -0.13 | 23 | 4.67E-23 | 0.69 | 0.82 | 3.63E-14 | 4.59E-28 | VULM01014500.1 | 149878 | 150204 |
| *INPP4B* | first intron | 0.10 | 10 | 1.65E-03 | 0.66 | 0.56 | 9.49E-10 | 2.89E-06 | VULM01010276.1 | 1726560 | 1726854 |
| *IRF1* | 3'-UTR | -0.10 | 13 | 2.36E-02 | 0.50 | 0.60 | 2.76E-07 | 1.22E-04 | VULM01005168.1 | 520964 | 521832 |
| *JPT1* | 3'-UTR | 0.11 | 10 | 5.46E-04 | 0.60 | 0.49 | 5.54E-13 | 6.40E-07 | VULM01001038.1 | 216690 | 217012 |
| *KCNA3* | 3'-UTR | -0.10 | 35 | 1.37E-13 | 0.31 | 0.41 | 4.17E-14 | 5.27E-18 | VULM01002219.1 | 82520 | 83501 |
| *KCNH5* | intron | 0.11 | 13 | 2.21E-04 | 0.61 | 0.50 | 4.28E-13 | 1.97E-07 | VULM01004113.1 | 1533355 | 1533979 |
| *KIF25* | 4757 bp downstream of TES | 0.11 | 10 | 8.30E-04 | 0.80 | 0.69 | 4.54E-10 | 1.13E-06 | VULM01006284.1 | 4748791 | 4749203 |
| *KLHL15* | first intron | 0.13 | 12 | 5.05E-06 | 0.62 | 0.48 | 4.30E-14 | 1.81E-09 | VULM01014277.1 | 55420 | 55551 |
| *KSR2* | 1583 bp downstream of TES | 0.16 | 10 | 1.86E-07 | 0.39 | 0.23 | 4.24E-14 | 3.43E-11 | VULM01008023.1 | 484831 | 484920 |
| *LAG3* | intron | 0.12 | 12 | 7.87E-04 | 0.42 | 0.30 | 4.09E-12 | 1.05E-06 | VULM01009205.1 | 427564 | 428468 |
| *LAMB3* | intron-exon boundary | -0.11 | 10 | 2.01E-04 | 0.42 | 0.54 | 1.26E-12 | 1.72E-07 | VULM01013663.1 | 1878475 | 1878778 |
| *LAMP1* | intron | -0.16 | 10 | 8.09E-04 | 0.43 | 0.59 | 4.27E-11 | 1.09E-06 | VULM01003239.1 | 2828734 | 2829166 |
| *LMNA* | first intron | 0.15 | 10 | 8.84E-10 | 0.53 | 0.38 | 4.31E-14 | 7.70E-14 | VULM01013242.1 | 133394 | 133479 |
| *LMO7* | intron | -0.12 | 10 | 4.60E-03 | 0.43 | 0.54 | 1.15E-11 | 1.18E-05 | VULM01011228.1 | 482583 | 482882 |
| *LRBA* | intron | -0.15 | 19 | 9.14E-11 | 0.32 | 0.47 | 4.42E-14 | 6.30E-15 | VULM01008730.1 | 1290376 | 1290583 |
| *LRCH3* | 377 bp downstream of TES | -0.17 | 12 | 1.02E-05 | 0.25 | 0.42 | 4.64E-14 | 4.28E-09 | VULM01004064.1 | 353983 | 354203 |
| *LUC7L* | 5'-UTR | 0.11 | 10 | 6.70E-04 | 0.40 | 0.29 | 2.09E-11 | 8.50E-07 | VULM01008195.1 | 401774 | 402001 |
| *MAMLD1* | intron | 0.18 | 10 | 1.97E-07 | 0.50 | 0.32 | 4.54E-14 | 3.69E-11 | VULM01009367.1 | 1255698 | 1255949 |
| *MAPKAP1* | intron | 0.12 | 10 | 5.86E-10 | 0.41 | 0.29 | 4.55E-14 | 4.73E-14 | VULM01013766.1 | 107587 | 107673 |
| *MBNL2* | 4968 bp downstream of TES | -0.11 | 10 | 8.94E-03 | 0.72 | 0.83 | 6.72E-11 | 2.99E-05 | VULM01002609.1 | 1296055 | 1296320 |
| *MDM2* | last exon to 3'-UTR | -0.15 | 10 | 4.10E-18 | 0.72 | 0.87 | 4.60E-14 | 7.21E-23 | VULM01012635.1 | 746748 | 747063 |
| *MDN1* | intron-exon boundary | -0.11 | 10 | 2.61E-02 | 0.49 | 0.59 | 5.15E-07 | 1.42E-04 | VULM01005939.1 | 4244079 | 4244962 |
| *MFNG* | 5'-UTR to first intron | 0.14 | 19 | 1.19E-08 | 0.48 | 0.34 | 4.25E-14 | 1.41E-12 | VULM01006140.1 | 518231 | 519061 |
| *MID2* | first intron | -0.10 | 10 | 7.50E-06 | 0.74 | 0.84 | 3.99E-14 | 2.94E-09 | VULM01003842.1 | 158518 | 158689 |
| *MID2* | intron | 0.12 | 13 | 3.22E-03 | 0.60 | 0.48 | 2.25E-13 | 7.25E-06 | VULM01003842.1 | 93443 | 93931 |
| *MIER1* | 974 bp upstream of TSS | -0.13 | 10 | 3.95E-03 | 0.54 | 0.67 | 4.97E-09 | 9.57E-06 | VULM01015833.1 | 1009909 | 1010329 |
| *MINDY4B* | 3'-UTR | -0.14 | 10 | 5.92E-04 | 0.37 | 0.51 | 3.12E-11 | 7.17E-07 | VULM01002742.1 | 1031497 | 1031728 |
| *MKNK2* | 1464 bp upstream of TSS | 0.10 | 10 | 9.64E-05 | 0.25 | 0.15 | 9.95E-14 | 6.79E-08 | VULM01009775.1 | 133092 | 133174 |
| *MLYCD* | first intron | 0.13 | 10 | 1.17E-02 | 0.68 | 0.55 | 1.21E-08 | 4.39E-05 | VULM01009081.1 | 975079 | 975808 |
| *MORN4* | 1176 bp upstream of TSS | 0.14 | 10 | 1.81E-03 | 0.42 | 0.28 | 1.91E-11 | 3.29E-06 | VULM01000731.1 | 42416 | 42578 |
| *MPDZ* | intron | -0.10 | 10 | 9.02E-03 | 0.52 | 0.63 | 2.72E-06 | 3.02E-05 | VULM01008671.1 | 951643 | 952793 |
| *MRGBP* | 4084 bp downstream of TES | -0.15 | 10 | 8.85E-05 | 0.39 | 0.54 | 1.04E-13 | 6.06E-08 | VULM01008304.1 | 1494754 | 1494863 |
| *MROH9* | intron | 0.18 | 10 | 1.25E-06 | 0.53 | 0.34 | 4.39E-14 | 3.35E-10 | VULM01014010.1 | 16181 | 16458 |
| *MRPL14* | 632 bp upstream of TSS | -0.12 | 10 | 7.56E-03 | 0.57 | 0.69 | 5.61E-09 | 2.34E-05 | VULM01006174.1 | 1704097 | 1705045 |
| *MRPS15* | 3'-UTR | 0.12 | 10 | 2.32E-03 | 0.46 | 0.34 | 6.05E-12 | 4.65E-06 | VULM01012774.1 | 89581 | 89736 |
| *MRPS15* | 3'-UTR | 0.19 | 12 | 1.50E-07 | 0.69 | 0.51 | 4.00E-14 | 2.65E-11 | VULM01012774.1 | 89787 | 89964 |
| *MYLK3* | 2449 bp upstream of TSS | 0.16 | 10 | 8.88E-06 | 0.76 | 0.60 | 4.23E-14 | 3.62E-09 | VULM01003799.1 | 3674874 | 3675958 |
| *MYO1D* | intron | 0.10 | 10 | 1.32E-04 | 0.87 | 0.77 | 1.05E-11 | 1.00E-07 | VULM01012770.1 | 91475 | 92465 |
| *MYO1D* | intron | -0.10 | 10 | 1.57E-05 | 0.77 | 0.87 | 4.39E-14 | 7.11E-09 | VULM01012770.1 | 109662 | 109851 |
| *MYO1D* | intron | -0.14 | 14 | 3.95E-03 | 0.41 | 0.55 | 2.82E-12 | 9.60E-06 | VULM01012770.1 | 116248 | 117390 |
| *MYO3B* | 570 bp downstream of TES | -0.10 | 10 | 2.14E-03 | 0.35 | 0.45 | 1.46E-12 | 4.16E-06 | VULM01009905.1 | 2331034 | 2331452 |
| *MYO5B* | intron | -0.17 | 12 | 2.42E-12 | 0.60 | 0.78 | 4.46E-14 | 1.19E-16 | VULM01007136.1 | 2063804 | 2064033 |
| *MYO9A* | intron | 0.12 | 13 | 3.79E-06 | 0.66 | 0.54 | 4.12E-14 | 1.23E-09 | VULM01008726.1 | 91313 | 91552 |
| *MYOC* | 3'-UTR | 0.12 | 10 | 4.87E-02 | 0.60 | 0.49 | 3.03E-08 | 3.56E-04 | VULM01014428.1 | 536636 | 536775 |
| *MYZAP* | 3480 bp downstream of TES | 0.13 | 10 | 6.09E-03 | 0.39 | 0.26 | 2.35E-06 | 1.73E-05 | VULM01004199.1 | 723088 | 723511 |
| *NAV1* | intron | -0.13 | 10 | 3.51E-04 | 0.46 | 0.59 | 6.71E-13 | 3.60E-07 | VULM01009778.1 | 57712 | 57968 |
| *NCMAP* | 560 bp upstream of TSS | 0.12 | 10 | 3.69E-02 | 0.62 | 0.50 | 1.67E-08 | 2.36E-04 | VULM01007190.1 | 225924 | 226117 |
| *NCOA7* | 5'-UTR | 0.12 | 10 | 2.42E-02 | 0.53 | 0.41 | 1.78E-10 | 1.27E-04 | VULM01006906.1 | 4691916 | 4692566 |
| *NCOR2* | intron | 0.11 | 12 | 8.20E-04 | 0.70 | 0.60 | 1.30E-11 | 1.11E-06 | VULM01011555.1 | 5286726 | 5287293 |
| *NCS1* | 5'-UTR to first intron | -0.12 | 10 | 1.54E-03 | 0.36 | 0.48 | 2.63E-10 | 2.64E-06 | VULM01010218.1 | 35207 | 35467 |
| *NDST2* | 5'-UTR | -0.11 | 10 | 2.28E-02 | 0.29 | 0.40 | 4.11E-09 | 1.16E-04 | VULM01004951.1 | 1364158 | 1364282 |
| *NET3* | first intron | 0.13 | 10 | 1.38E-03 | 0.55 | 0.41 | 1.33E-11 | 2.26E-06 | VULM01006817.1 | 421849 | 422056 |
| *NFASC* | intron-exon boundary | -0.11 | 10 | 4.84E-04 | 0.37 | 0.48 | 1.17E-07 | 5.50E-07 | VULM01013663.1 | 181967 | 182329 |
| *NFXL1* | 341 bp downstream of TES | -0.10 | 11 | 1.35E-03 | 0.71 | 0.81 | 4.65E-12 | 2.18E-06 | VULM01014555.1 | 682400 | 683290 |
| *NIN* | intron | 0.16 | 25 | 1.80E-15 | 0.63 | 0.47 | 4.79E-14 | 4.44E-20 | VULM01000525.1 | 253660 | 254334 |
| *NOS1AP* | 3115 bp upstream of TSS | -0.10 | 10 | 5.36E-04 | 0.37 | 0.47 | 9.02E-13 | 6.25E-07 | VULM01015885.1 | 1035329 | 1035487 |
| *NR0B2* | first intron | -0.12 | 10 | 4.39E-03 | 0.38 | 0.50 | 1.96E-11 | 1.10E-05 | VULM01003975.1 | 1883 | 2001 |
| *NR3C1* | 1351 bp upstream of TSS | -0.12 | 10 | 1.52E-02 | 0.32 | 0.44 | 6.63E-11 | 6.41E-05 | VULM01005168.1 | 1403879 | 1403960 |
| *NRK* | 4351 bp upstream of TSS | 0.12 | 10 | 3.63E-04 | 0.76 | 0.63 | 4.90E-14 | 3.78E-07 | VULM01009367.1 | 2058996 | 2059124 |
| *NRTN* | 2137 bp upstream of TSS | 0.12 | 10 | 2.16E-02 | 0.57 | 0.45 | 1.17E-08 | 1.08E-04 | VULM01013795.1 | 210598 | 210708 |
| *NRXN1* | intron-exon boundary | 0.16 | 10 | 7.30E-06 | 0.57 | 0.41 | 4.62E-14 | 2.85E-09 | VULM01006330.1 | 3325306 | 3325612 |
| *NSMF* | 3'-UTR | 0.12 | 10 | 1.18E-04 | 0.63 | 0.50 | 4.27E-13 | 8.69E-08 | VULM01007130.1 | 866342 | 866429 |
| *NT5E* | intron | 0.16 | 15 | 9.29E-13 | 0.88 | 0.71 | 4.57E-14 | 4.31E-17 | VULM01003851.1 | 1182535 | 1183280 |
| *PACS2* | intron | 0.11 | 12 | 1.15E-02 | 0.48 | 0.38 | 1.31E-06 | 4.28E-05 | VULM01008931.1 | 1179424 | 1180205 |
| *PAG1* | 3'-UTR | -0.11 | 10 | 6.70E-03 | 0.48 | 0.59 | 2.30E-08 | 1.97E-05 | VULM01014069.1 | 932998 | 934122 |
| *PARD6A* | first intron | -0.12 | 13 | 7.60E-07 | 0.56 | 0.68 | 4.20E-14 | 1.88E-10 | VULM01010374.1 | 607750 | 608000 |
| *PASD1* | intron | -0.12 | 10 | 2.44E-03 | 0.56 | 0.68 | 4.42E-09 | 5.01E-06 | VULM01009367.1 | 1793223 | 1793804 |
| *PCSK7* | intron | -0.12 | 11 | 2.97E-02 | 0.45 | 0.57 | 9.97E-09 | 1.71E-04 | VULM01002389.1 | 133967 | 134198 |
| *PDE7B* | intron | -0.12 | 10 | 1.25E-06 | 0.39 | 0.51 | 2.39E-08 | 3.37E-10 | VULM01007568.1 | 553627 | 554283 |
| *PDE8B* | intron-exon boundary | -0.18 | 12 | 9.14E-11 | 0.34 | 0.52 | 4.20E-14 | 6.35E-15 | VULM01006363.1 | 1698045 | 1698297 |
| *PDGFRA* | 1867 bp upstream of TSS | 0.10 | 12 | 1.09E-05 | 0.61 | 0.51 | 4.26E-14 | 4.62E-09 | VULM01014642.1 | 518029 | 518143 |
| *PDZK1* | 5'-UTR | -0.11 | 10 | 2.58E-04 | 0.75 | 0.85 | 3.70E-13 | 2.39E-07 | VULM01013471.1 | 12577477 | 12578025 |
| *PERP* | 103 bp upstream of TSS | 0.13 | 10 | 2.50E-03 | 0.47 | 0.35 | 2.04E-08 | 5.17E-06 | VULM01006140.1 | 1009263 | 1009599 |
| *PHACTR4* | intron | 0.10 | 10 | 5.38E-06 | 0.76 | 0.66 | 1.58E-12 | 1.94E-09 | VULM01012527.1 | 29757 | 30146 |
| *PKIA* | 640 bp upstream of TSS | -0.10 | 10 | 2.17E-03 | 0.53 | 0.64 | 6.91E-07 | 4.25E-06 | VULM01014069.1 | 1914689 | 1916043 |
| *PLPP3* | intron | -0.11 | 10 | 1.91E-03 | 0.27 | 0.37 | 3.76E-08 | 3.54E-06 | VULM01011261.1 | 1572593 | 1572739 |
| *PLPP3* | intron | -0.11 | 10 | 3.61E-04 | 0.46 | 0.57 | 2.86E-07 | 3.76E-07 | VULM01011261.1 | 1585378 | 1585741 |
| *PLXNC1* | 5'-UTR | 0.11 | 39 | 6.11E-06 | 0.53 | 0.42 | 4.06E-14 | 2.28E-09 | VULM01000246.1 | 3274460 | 3275062 |
| *PMP22* | intron | -0.12 | 10 | 2.36E-03 | 0.51 | 0.64 | 1.97E-09 | 4.79E-06 | VULM01002161.1 | 204272 | 204746 |
| *POC1A* | 1531 bp upstream of TSS | 0.14 | 10 | 1.40E-06 | 0.62 | 0.48 | 1.84E-13 | 3.88E-10 | VULM01013099.1 | 267294 | 267607 |
| *PPP1R12B* | intron-exon boundary | 0.10 | 10 | 4.28E-08 | 0.37 | 0.27 | 9.70E-14 | 5.98E-12 | VULM01014342.1 | 56661 | 57008 |
| *PPP1R9A* | 5'-UTR | 0.11 | 10 | 1.09E-03 | 0.59 | 0.48 | 1.91E-10 | 1.60E-06 | VULM01006696.1 | 808966 | 809340 |
| *PPP4R2* | 502 bp downstream of TES | -0.11 | 16 | 2.39E-04 | 0.49 | 0.60 | 3.49E-13 | 2.18E-07 | VULM01009947.1 | 3946634 | 3947367 |
| *PRELID2* | intron | -0.16 | 10 | 6.98E-11 | 0.69 | 0.86 | 4.47E-14 | 4.51E-15 | VULM01005168.1 | 1954612 | 1955584 |
| *PSD3* | first intron | -0.12 | 12 | 9.96E-07 | 0.24 | 0.36 | 6.36E-14 | 2.57E-10 | VULM01002801.1 | 243580 | 243761 |
| *PSTPIP1* | first intron | 0.13 | 10 | 2.62E-03 | 0.67 | 0.54 | 5.24E-11 | 5.49E-06 | VULM01008526.1 | 545772 | 546716 |
| *PUS10* | intron-exon boundary | -0.13 | 10 | 3.71E-04 | 0.68 | 0.82 | 1.17E-13 | 3.89E-07 | VULM01013711.1 | 355839 | 356180 |
| *R3HDM4* | intron | -0.15 | 10 | 1.13E-07 | 0.38 | 0.52 | 4.54E-14 | 1.84E-11 | VULM01010829.1 | 6161 | 6350 |
| *RAB27A* | 491 bp upstream of TSS | -0.20 | 15 | 5.11E-13 | 0.35 | 0.55 | 3.97E-14 | 2.20E-17 | VULM01011934.1 | 2381328 | 2381577 |
| *RAB3B* | first intron | 0.11 | 12 | 1.82E-03 | 0.65 | 0.54 | 9.34E-10 | 3.32E-06 | VULM01002583.1 | 122238 | 122568 |
| *RAI2* | 5'-UTR | -0.16 | 10 | 4.50E-08 | 0.45 | 0.60 | 4.27E-14 | 6.36E-12 | VULM01004673.1 | 2755065 | 2755293 |
| *RAPGEF5* | intron | -0.13 | 10 | 4.67E-04 | 0.48 | 0.62 | 3.87E-12 | 5.23E-07 | VULM01002496.1 | 560900 | 561224 |
| *RASSF5* | first intron | 0.12 | 26 | 1.10E-11 | 0.48 | 0.36 | 4.05E-14 | 6.36E-16 | VULM01013663.1 | 962394 | 962812 |
| *RBMS3* | first intron | 0.10 | 12 | 4.41E-07 | 0.52 | 0.42 | 4.66E-14 | 1.00E-10 | VULM01002123.1 | 1789185 | 1790072 |
| *REL* | intron | -0.12 | 20 | 2.26E-06 | 0.39 | 0.51 | 4.27E-14 | 6.73E-10 | VULM01013711.1 | 374180 | 374568 |
| *RGP1* | intron | 0.10 | 10 | 2.76E-02 | 0.56 | 0.45 | 1.11E-07 | 1.54E-04 | VULM01009490.1 | 91937 | 92058 |
| *RLIM* | 3'-UTR | 0.10 | 10 | 1.28E-07 | 0.65 | 0.55 | 4.33E-14 | 2.16E-11 | VULM01010275.1 | 4939915 | 4940069 |
| *RNASEH2B* | 621 bp upstream of TSS | 0.13 | 12 | 4.32E-05 | 0.49 | 0.36 | 5.56E-10 | 2.41E-08 | VULM01005679.1 | 261051 | 261510 |
| *RNF220* | first intron | -0.12 | 10 | 2.49E-02 | 0.47 | 0.59 | 4.03E-08 | 1.32E-04 | VULM01014864.1 | 313575 | 314124 |
| *RUNX2* | intron | 0.11 | 10 | 3.26E-02 | 0.70 | 0.59 | 1.04E-07 | 1.96E-04 | VULM01010451.1 | 1211972 | 1212167 |
| *SAMD3* | intron | 0.11 | 10 | 6.40E-07 | 0.73 | 0.63 | 4.66E-14 | 1.55E-10 | VULM01006906.1 | 2496570 | 2496812 |
| *SARM1* | intron-exon boundary | -0.15 | 32 | 2.69E-24 | 0.60 | 0.75 | 4.75E-14 | 2.26E-29 | VULM01003115.1 | 112025 | 113302 |
| *SCFD1* | 181 bp downstream of TES | 0.11 | 10 | 6.73E-03 | 0.80 | 0.68 | 1.01E-09 | 1.98E-05 | VULM01010130.1 | 784709 | 785335 |
| *SEPTIN9* | intron | 0.11 | 10 | 6.20E-04 | 0.66 | 0.56 | 3.86E-11 | 7.64E-07 | VULM01008732.1 | 724819 | 725252 |
| *SFTPA1* | 3284 bp upstream of TSS | 0.12 | 10 | 2.58E-02 | 0.70 | 0.58 | 3.97E-09 | 1.40E-04 | VULM01006019.1 | 3694053 | 3694741 |
| *SH3GL1* | first intron | 0.13 | 10 | 8.84E-04 | 0.73 | 0.60 | 8.84E-12 | 1.22E-06 | VULM01002240.1 | 12273 | 12375 |
| *SH3GL1* | first intron | 0.13 | 20 | 6.37E-05 | 0.59 | 0.46 | 4.74E-14 | 3.95E-08 | VULM01002240.1 | 12647 | 13079 |
| *SH3GL1* | first intron | 0.20 | 10 | 2.31E-07 | 0.72 | 0.52 | 3.95E-14 | 4.47E-11 | VULM01002240.1 | 13090 | 13300 |
| *SH3GL1* | first intron | 0.14 | 13 | 1.16E-05 | 0.64 | 0.51 | 4.39E-14 | 5.00E-09 | VULM01002240.1 | 13376 | 13536 |
| *SH3KBP1* | first intron | -0.10 | 10 | 2.36E-02 | 0.54 | 0.64 | 3.25E-06 | 1.23E-04 | VULM01004673.1 | 1864732 | 1865346 |
| *SHLD2* | 3706 bp downstream of TES | 0.16 | 15 | 2.72E-11 | 0.51 | 0.35 | 4.10E-14 | 1.68E-15 | VULM01006019.1 | 5738666 | 5738988 |
| *SHROOM3* | 3'-UTR | -0.17 | 10 | 7.75E-08 | 0.27 | 0.44 | 4.05E-14 | 1.18E-11 | VULM01003541.1 | 449455 | 449772 |
| *SIGIRR* | 5'-UTR | 0.10 | 10 | 4.86E-03 | 0.65 | 0.54 | 2.11E-09 | 1.27E-05 | VULM01014905.1 | 1416 | 1663 |
| *SIK3* | first intron | -0.10 | 17 | 1.58E-06 | 0.66 | 0.76 | 4.23E-14 | 4.47E-10 | VULM01002070.1 | 51835 | 52160 |
| *SIL1* | 5'-UTR | -0.14 | 19 | 1.48E-16 | 0.71 | 0.85 | 3.89E-14 | 3.32E-21 | VULM01006763.1 | 7531279 | 7531587 |
| *SIVA* | 521 bp downstream of TES | -0.11 | 13 | 2.33E-07 | 0.56 | 0.67 | 6.69E-14 | 4.53E-11 | VULM01000108.1 | 1091629 | 1093582 |
| *SLC23A2* | 2448 bp downstream of TES | -0.10 | 12 | 1.53E-02 | 0.30 | 0.41 | 2.70E-08 | 6.47E-05 | VULM01012755.1 | 647 | 827 |
| *SLC26A8* | intron-exon boundary | -0.11 | 13 | 2.66E-03 | 0.61 | 0.72 | 1.90E-09 | 5.63E-06 | VULM01012884.1 | 135636 | 136440 |
| *SLC38A3* | intron | -0.13 | 15 | 7.17E-08 | 0.53 | 0.66 | 4.64E-14 | 1.07E-11 | VULM01010865.1 | 45544 | 46012 |
| *SLC66A2* | 3'-UTR | -0.11 | 10 | 1.25E-02 | 0.61 | 0.72 | 2.51E-08 | 4.85E-05 | VULM01007120.1 | 694695 | 695176 |
| *SLC6A13* | intron | -0.16 | 13 | 1.52E-04 | 0.57 | 0.73 | 4.16E-14 | 1.22E-07 | VULM01001605.1 | 85600 | 85864 |
| *SLIT3* | intron | 0.13 | 12 | 1.24E-09 | 0.80 | 0.67 | 4.47E-14 | 1.13E-13 | VULM01006763.1 | 4769500 | 4769805 |
| *SLX4* | intron-exon boundary | -0.13 | 10 | 2.16E-03 | 0.69 | 0.81 | 1.95E-13 | 4.21E-06 | VULM01002535.1 | 10694 | 10989 |
| *SNAPC4* | intron | 0.10 | 12 | 1.30E-04 | 0.76 | 0.66 | 3.13E-13 | 9.79E-08 | VULM01003774.1 | 849281 | 849433 |

| | | | | | | | | | | | |
|---|---|---|---|---|---|---|---|---|---|---|---|
| *SNF2L2* | 642 bp upstream of TSS | -0.15 | 12 | 4.81E-12 | 0.41 | 0.56 | 4.07E-14 | 2.57E-16 | VULM01000844.1 | 869135 | 870293 |
| *SPIDR* | intron | -0.11 | 10 | 2.13E-06 | 0.14 | 0.25 | 4.10E-14 | 6.27E-10 | VULM01002927.1 | 3113979 | 3114272 |
| *SPINK4* | 3242 bp upstream of TSS | 0.10 | 10 | 2.09E-04 | 0.34 | 0.23 | 1.35E-07 | 1.84E-07 | VULM01008176.1 | 498896 | 499188 |
| *SSBP2* | 1223 bp upstream of TSS | -0.19 | 10 | 5.98E-10 | 0.27 | 0.47 | 4.59E-14 | 4.91E-14 | VULM01007843.1 | 786004 | 786169 |
| *SSH1* | 3'-UTR | -0.11 | 10 | 3.00E-03 | 0.40 | 0.51 | 1.69E-07 | 6.62E-06 | VULM01000519.1 | 9617 | 9857 |
| *STAB2* | intron-exon boundary | 0.10 | 10 | 1.20E-02 | 0.51 | 0.41 | 2.90E-06 | 4.54E-05 | VULM01002643.1 | 886991 | 887439 |
| *STRN* | intron | -0.12 | 10 | 6.82E-03 | 0.63 | 0.75 | 6.03E-11 | 2.02E-05 | VULM01012098.1 | 354522 | 355979 |
| *TBL1X* | 5'-UTR | 0.13 | 10 | 1.10E-02 | 0.70 | 0.56 | 3.25E-11 | 3.97E-05 | VULM01015136.1 | 2393782 | 2394059 |
| *TCF4* | intron | 0.10 | 10 | 8.62E-04 | 0.60 | 0.50 | 1.01E-13 | 1.18E-06 | VULM01007136.1 | 330447 | 330591 |
| *TEF* | 3'-UTR | 0.12 | 10 | 1.41E-03 | 0.83 | 0.72 | 1.13E-12 | 2.33E-06 | VULM01008453.1 | 541447 | 542130 |
| *TJAP1* | 1284 bp downstream of TES | -0.12 | 17 | 2.05E-07 | 0.60 | 0.72 | 4.61E-14 | 3.90E-11 | VULM01006174.1 | 1044624 | 1044968 |
| *TLE4* | first intron | -0.10 | 12 | 8.38E-04 | 0.25 | 0.36 | 6.48E-10 | 1.14E-06 | VULM01005592.1 | 242881 | 243197 |
| *TMCC3* | intron-exon boundary | -0.16 | 11 | 1.23E-06 | 0.17 | 0.33 | 1.08E-13 | 3.28E-10 | VULM01000246.1 | 3159423 | 3160134 |
| *TMEM45B* | 957 bp downstream of TES | 0.10 | 10 | 3.99E-04 | 0.79 | 0.69 | 3.80E-07 | 4.23E-07 | VULM01008572.1 | 329467 | 329795 |
| *TMPRSS13* | 1540 bp downstream of TES | 0.18 | 10 | 3.87E-05 | 0.61 | 0.43 | 4.34E-14 | 2.09E-08 | VULM01009713.1 | 175083 | 175231 |
| *TMTC1* | intron | -0.11 | 10 | 3.93E-03 | 0.17 | 0.27 | 9.95E-11 | 9.53E-06 | VULM01001605.1 | 603247 | 603808 |
| *TNNC1* | intron-exon boundary | -0.11 | 13 | 7.73E-05 | 0.55 | 0.66 | 1.02E-11 | 5.11E-08 | VULM01013099.1 | 758226 | 758510 |
| *TNS4* | 2648 bp downstream of TES | -0.11 | 10 | 2.41E-02 | 0.39 | 0.50 | 9.79E-09 | 1.27E-04 | VULM01007974.1 | 240469 | 240712 |
| *TPRA1* | 525 bp upstream of TSS | -0.11 | 12 | 1.19E-04 | 0.58 | 0.68 | 4.82E-14 | 8.83E-08 | VULM01015089.1 | 229376 | 229522 |
| *TRABD2B* | first intron | -0.12 | 10 | 2.59E-03 | 0.34 | 0.46 | 4.81E-12 | 5.43E-06 | VULM01002583.1 | 2310602 | 2311133 |
| *TRHR* | 3183 bp upstream of TSS | -0.16 | 10 | 1.21E-09 | 0.49 | 0.65 | 4.32E-14 | 1.09E-13 | VULM01005733.1 | 62083 | 62290 |
| *TRIM13* | 6024 bp downstream of TES | 0.13 | 19 | 1.09E-22 | 0.25 | 0.12 | 3.86E-14 | 1.23E-27 | VULM01005679.1 | 803370 | 803719 |
| *TRPV2* | first intron | 0.11 | 10 | 2.13E-05 | 0.64 | 0.53 | 4.07E-10 | 1.03E-08 | VULM01000885.1 | 204993 | 205761 |
| *TTC7A* | 2698 bp downstream of TES | -0.16 | 15 | 1.01E-07 | 0.43 | 0.59 | 4.33E-14 | 1.61E-11 | VULM01006174.1 | 5479537 | 5479955 |
| *TTYH3* | first intron | 0.14 | 10 | 1.76E-06 | 0.51 | 0.36 | 4.32E-14 | 5.04E-10 | VULM01014983.1 | 1112101 | 1112324 |
| *TWIST2* | 5869 bp upstream of TSS | -0.18 | 15 | 7.19E-27 | 0.20 | 0.39 | 4.07E-14 | 4.54E-32 | VULM01000284.1 | 84369 | 84544 |
| *TXNDC16* | intron | -0.10 | 10 | 3.15E-02 | 0.56 | 0.67 | 7.26E-08 | 1.85E-04 | VULM01007008.1 | 22681 | 23048 |
| *TXNDC16* | last exon to 3'-UTR | 0.18 | 13 | 4.81E-11 | 0.51 | 0.32 | 4.56E-14 | 3.04E-15 | VULM01007008.1 | 19273 | 19882 |
| *U2SURP* | first exon | 0.11 | 16 | 1.86E-08 | 0.34 | 0.23 | 4.39E-14 | 2.33E-12 | VULM01005154.1 | 1496769 | 1496973 |
| *UBL3* | 4396 bp downstream of TES | 0.10 | 10 | 3.16E-06 | 0.58 | 0.48 | 4.60E-14 | 9.89E-10 | VULM01010319.1 | 5494427 | 5494657 |
| *UGGT2* | 4737 bp downstream of TES | -0.15 | 12 | 1.02E-10 | 0.41 | 0.56 | 4.06E-14 | 7.14E-15 | VULM01002609.1 | 1901307 | 1901657 |
| *UNC50* | 3'-UTR | -0.12 | 10 | 1.04E-02 | 0.43 | 0.55 | 8.12E-09 | 3.72E-05 | VULM01007759.1 | 5941386 | 5941855 |
| *UPF2* | 3791 bp downstream of TES | 0.11 | 13 | 9.53E-09 | 0.38 | 0.27 | 4.30E-14 | 1.11E-12 | VULM01006483.1 | 1904146 | 1904243 |
| *USH1G* | 5'-UTR | 0.12 | 16 | 7.30E-05 | 0.74 | 0.63 | 4.49E-14 | 4.73E-08 | VULM01001038.1 | 120039 | 120697 |
| *USP12* | 5'-UTR | 0.13 | 10 | 1.39E-02 | 0.67 | 0.54 | 8.06E-08 | 5.62E-05 | VULM01010275.1 | 5090357 | 5090684 |
| *USP18* | 654 bp downstream of TES | 0.13 | 10 | 1.48E-04 | 0.61 | 0.48 | 1.29E-13 | 1.17E-07 | VULM01001894.1 | 2970907 | 2971571 |
| *USP28* | 2238 bp upstream of TSS | 0.12 | 10 | 4.97E-06 | 0.81 | 0.69 | 4.67E-14 | 1.77E-09 | VULM01013870.1 | 13303 | 13453 |
| *USP28* | 1565 bp upstream of TSS | 0.16 | 12 | 9.14E-11 | 0.70 | 0.54 | 4.52E-14 | 6.27E-15 | VULM01013870.1 | 13701 | 14126 |
| *USP28* | 975 bp upstream of TSS | 0.10 | 15 | 1.72E-04 | 0.73 | 0.63 | 4.63E-14 | 1.42E-07 | VULM01013870.1 | 14491 | 14716 |
| *USP28* | first intron | -0.12 | 10 | 1.38E-04 | 0.59 | 0.71 | 1.20E-12 | 1.06E-07 | VULM01013870.1 | 16453 | 16622 |
| *USP7* | 5'-UTR | -0.14 | 10 | 8.24E-06 | 0.45 | 0.59 | 4.52E-14 | 3.29E-09 | VULM01007313.1 | 2491338 | 2491885 |
| *UST* | intron | 0.12 | 10 | 3.42E-07 | 0.38 | 0.26 | 2.76E-12 | 7.09E-11 | VULM01006284.1 | 11747456 | 11747582 |
| *VAC14* | intron | -0.11 | 10 | 6.77E-03 | 0.65 | 0.77 | 3.61E-12 | 2.00E-05 | VULM01008312.1 | 10638 | 10818 |
| *VAT1L* | intron | -0.13 | 10 | 7.72E-07 | 0.65 | 0.77 | 4.20E-14 | 1.92E-10 | VULM01014523.1 | 1598376 | 1598954 |
| *VPS13B* | intron | 0.10 | 10 | 7.39E-07 | 0.70 | 0.60 | 4.55E-14 | 1.82E-10 | VULM01004535.1 | 252459 | 252619 |
| *VPS13C* | intron-exon boundary | 0.17 | 10 | 3.04E-05 | 0.37 | 0.21 | 4.90E-14 | 1.56E-08 | VULM01005615.1 | 829810 | 830274 |
| *VWF* | intron | 0.12 | 10 | 1.79E-03 | 0.74 | 0.63 | 1.52E-11 | 3.23E-06 | VULM01008758.1 | 1644218 | 1645345 |
| *WDR31* | exon | 0.10 | 10 | 1.25E-02 | 0.66 | 0.56 | 1.82E-10 | 4.84E-05 | VULM01007130.1 | 1225875 | 1225995 |
| *WDR76* | 3378 bp downstream of TES | 0.17 | 22 | 1.46E-20 | 0.38 | 0.22 | 4.34E-14 | 2.25E-25 | VULM01008304.1 | 2412751 | 2413025 |
| *WFDC2* | first intron | -0.10 | 37 | 1.07E-25 | 0.56 | 0.66 | 5.57E-14 | 8.27E-31 | VULM01000812.1 | 6203 | 6598 |
| *WNT5B* | intron | 0.10 | 10 | 2.39E-02 | 0.63 | 0.53 | 2.14E-06 | 1.25E-04 | VULM01001894.1 | 4297064 | 4297217 |
| *WSCD2* | 173 bp downstream of TES | 0.10 | 10 | 6.01E-04 | 0.46 | 0.35 | 3.59E-11 | 7.32E-07 | VULM01005923.1 | 814917 | 815135 |
| *WWOX* | intron | -0.10 | 10 | 2.42E-03 | 0.32 | 0.42 | 3.81E-10 | 4.94E-06 | VULM01014523.1 | 1233148 | 1233554 |
| *ZBTB16* | first intron | -0.10 | 10 | 3.32E-02 | 0.61 | 0.71 | 9.31E-09 | 2.01E-04 | VULM01015150.1 | 43365 | 43632 |
| *ZBTB16* | first intron | -0.10 | 10 | 2.62E-03 | 0.17 | 0.28 | 1.62E-11 | 5.49E-06 | VULM01015150.1 | 73742 | 73875 |
| *ZC3H3* | intron | 0.12 | 10 | 1.98E-02 | 0.57 | 0.46 | 5.85E-07 | 9.41E-05 | VULM01001519.1 | 3266089 | 3266549 |
| *ZC3H3* | intron | 0.15 | 10 | 2.10E-05 | 0.42 | 0.27 | 4.69E-14 | 1.01E-08 | VULM01001519.1 | 3267131 | 3267426 |
| *ZDHHC1* | 2040 bp downstream of TES | 0.11 | 22 | 2.16E-08 | 0.37 | 0.26 | 4.30E-14 | 2.75E-12 | VULM01006027.1 | 248079 | 248374 |
| *ZGRF1* | intron-exon boundary | -0.18 | 10 | 2.39E-06 | 0.38 | 0.57 | 5.38E-13 | 7.21E-10 | VULM01008832.1 | 12870 | 13332 |
| *ZHX1* | 3'-UTR | 0.10 | 13 | 1.04E-03 | 0.65 | 0.54 | 4.67E-12 | 1.51E-06 | VULM01004441.1 | 1007242 | 1007629 |
| *ZMIZ1* | 5'-UTR | 0.11 | 10 | 3.50E-03 | 0.63 | 0.52 | 2.58E-10 | 8.12E-06 | VULM01006019.1 | 2336040 | 2336392 |
| *ZNF512B* | intron | -0.14 | 10 | 5.56E-04 | 0.42 | 0.56 | 1.55E-11 | 6.58E-07 | VULM01014731.1 | 9523 | 9761 |
| *ZNF512B* | intron | 0.17 | 14 | 1.23E-10 | 0.45 | 0.28 | 4.01E-14 | 8.91E-15 | VULM01014731.1 | 24627 | 24950 |
| *ZNF536* | intron | 0.11 | 14 | 1.47E-09 | 0.48 | 0.37 | 4.30E-14 | 1.39E-13 | VULM01003799.1 | 2243663 | 2243757 |
| *ZNF541* | intron | -0.12 | 32 | 3.67E-21 | 0.59 | 0.71 | 4.76E-14 | 4.64E-26 | VULM01008151.1 | 9464 | 9863 |
| *ZNF850* | 3'-UTR | -0.10 | 10 | 2.62E-03 | 0.61 | 0.71 | 1.36E-09 | 5.50E-06 | VULM01009736.1 | 455181 | 455402 |
| *(predicted orf)* | 4351 bp upstream of TSS | 0.13 | 10 | 1.39E-02 | 0.67 | 0.54 | 8.06E-08 | 5.62E-05 | VULM01010275.1 | 5090357 | 5090684 |
| *(predicted orf)* | 2266 bp upstream of TSS | -0.12 | 10 | 1.03E-03 | 0.51 | 0.63 | 1.04E-08 | 1.50E-06 | VULM01008572.1 | 225882 | 226078 |
| *(predicted orf)* | 2172 bp upstream of TSS | -0.14 | 10 | 8.24E-06 | 0.45 | 0.59 | 4.52E-14 | 3.29E-09 | VULM01007313.1 | 2491338 | 2491885 |
| *(predicted orf)* | 1774 bp upstream of TSS | 0.11 | 10 | 6.70E-04 | 0.40 | 0.29 | 2.09E-11 | 8.50E-07 | VULM01008195.1 | 401774 | 402001 |
| *(predicted orf)* | 1561 bp upstream of TSS | -0.12 | 10 | 3.47E-03 | 0.62 | 0.73 | 3.71E-10 | 8.01E-06 | VULM01012584.1 | 24981 | 25133 |
| *(predicted orf)* | 1103 bp upstream of TSS | 0.10 | 10 | 4.81E-02 | 0.66 | 0.56 | 7.05E-08 | 3.50E-04 | VULM01015833.1 | 895744 | 896086 |
| *(predicted orf)* | 835 bp upstream of TSS | -0.10 | 10 | 5.29E-04 | 0.70 | 0.81 | 1.18E-13 | 6.12E-07 | VULM01010276.1 | 1745129 | 1745838 |
| *(predicted orf)* | 141 bp upstream of TSS | 0.19 | 15 | 1.17E-31 | 0.88 | 0.69 | 5.26E-14 | 3.30E-37 | VULM01001894.1 | 2103949 | 2104323 |
| *(predicted orf)* | 94 bp upstream of TSS | -0.11 | 10 | 1.61E-02 | 0.26 | 0.37 | 9.92E-08 | 6.91E-05 | VULM01010374.1 | 654685 | 654906 |
| *(predicted orf)* | first intron | 0.11 | 10 | 2.13E-05 | 0.64 | 0.53 | 4.07E-10 | 1.03E-08 | VULM01000885.1 | 204993 | 205761 |
| *(predicted orf)* | first intron | 0.11 | 10 | 1.48E-03 | 0.55 | 0.44 | 1.92E-07 | 2.48E-06 | VULM01003306.1 | 4855809 | 4856431 |
| *(predicted orf)* | first intron | 0.12 | 16 | 3.07E-07 | 0.84 | 0.72 | 4.35E-14 | 6.26E-11 | VULM01003719.1 | 226582 | 227157 |
| *(predicted orf)* | first intron | -0.12 | 13 | 7.60E-07 | 0.56 | 0.68 | 4.20E-14 | 1.88E-10 | VULM01010374.1 | 607750 | 608000 |
| *(predicted orf)* | first intron | -0.11 | 13 | 1.52E-03 | 0.42 | 0.52 | 8.64E-10 | 2.58E-06 | VULM01003902.1 | 1159449 | 1159708 |
| *(predicted orf)* | first intron | -0.13 | 13 | 6.80E-04 | 0.47 | 0.60 | 4.31E-14 | 8.70E-07 | VULM01015654.1 | 81786 | 82053 |
| *(predicted orf)* | intron | 0.10 | 12 | 1.30E-04 | 0.76 | 0.66 | 3.13E-13 | 9.79E-08 | VULM01003774.1 | 849281 | 849433 |
| *(predicted orf)* | intron | 0.11 | 10 | 4.85E-06 | 0.51 | 0.40 | 7.86E-13 | 1.72E-09 | VULM01005385.1 | 29903 | 30269 |
| *(predicted orf)* | intron | -0.13 | 10 | 5.33E-04 | 0.58 | 0.71 | 9.13E-14 | 6.20E-07 | VULM01008382.1 | 1365241 | 1365423 |
| *(predicted orf)* | intron | 0.25 | 16 | 8.05E-21 | 0.76 | 0.51 | 4.59E-14 | 1.07E-25 | VULM01015654.1 | 48059 | 48552 |
| *(predicted orf)* | exon | -0.14 | 10 | 4.18E-09 | 0.70 | 0.84 | 4.66E-14 | 4.46E-13 | VULM01003902.1 | 1159361 | 1159426 |
| *(predicted orf)* | exon | -0.16 | 12 | 1.93E-15 | 0.50 | 0.66 | 4.11E-14 | 4.87E-20 | VULM01015837.1 | 214914 | 215067 |
| *(predicted orf)* | last exon to 3'-UTR | 0.11 | 16 | 1.86E-08 | 0.34 | 0.23 | 4.39E-14 | 2.33E-12 | VULM01005154.1 | 1496769 | 1496973 |
| *(predicted orf)* | last intron to 3'-UTR | -0.23 | 21 | 5.18E-23 | 0.30 | 0.53 | 4.13E-14 | 5.45E-28 | VULM01003115.1 | 111433 | 112023 |
| *(predicted orf)* | last intron to 3'-UTR | 0.14 | 10 | 3.92E-02 | 0.40 | 0.26 | 4.16E-08 | 2.58E-04 | VULM01014838.1 | 1058080 | 1058567 |
| *(predicted orf)* | last intron to 3'-UTR | 0.19 | 10 | 7.14E-06 | 0.62 | 0.42 | 5.41E-14 | 2.77E-09 | VULM01014838.1 | 1058597 | 1059350 |
| *(predicted orf)* | 3'-UTR | -0.10 | 10 | 1.18E-02 | 0.49 | 0.60 | 7.38E-06 | 4.46E-05 | VULM01000184.1 | 137479 | 138092 |
| *(predicted orf)* | 3'-UTR | -0.19 | 10 | 8.84E-14 | 0.51 | 0.70 | 3.71E-14 | 3.29E-18 | VULM01010856.1 | 1137476 | 1137640 |
| *(predicted orf)* | 5'-UTR | 0.10 | 10 | 3.06E-02 | 0.49 | 0.39 | 1.07E-05 | 1.79E-04 | VULM01000128.1 | 2481315 | 2481511 |
| *(predicted orf)* | 102 bp downstream of TES | -0.10 | 13 | 2.36E-02 | 0.50 | 0.60 | 2.76E-07 | 1.22E-04 | VULM01005168.1 | 520964 | 521832 |
| *(predicted orf)* | 211 bp downstream of TES | -0.14 | 10 | 1.46E-04 | 0.49 | 0.63 | 8.35E-14 | 1.15E-07 | VULM01005296.1 | 1151175 | 1151263 |
| *(predicted orf)* | 211 bp downstream of TES | -0.12 | 10 | 1.93E-03 | 0.46 | 0.58 | 5.41E-11 | 3.61E-06 | VULM01009358.1 | 150795 | 151179 |
| *(predicted orf)* | 313 bp downstream of TES | -0.13 | 12 | 3.18E-06 | 0.68 | 0.81 | 4.61E-14 | 9.98E-10 | VULM01004759.1 | 68201 | 68556 |
| *(predicted orf)* | 377 bp downstream of TES | -0.17 | 12 | 1.02E-05 | 0.25 | 0.42 | 4.64E-14 | 4.28E-09 | VULM01004064.1 | 353983 | 354203 |
| *(predicted orf)* | 816 bp downstream of TES | 0.13 | 17 | 3.45E-11 | 0.49 | 0.36 | 4.29E-14 | 2.16E-15 | VULM01012755.1 | 113486 | 113793 |
| *(predicted orf)* | 974 bp downstream of TES | -0.13 | 10 | 3.95E-03 | 0.54 | 0.67 | 4.97E-09 | 9.57E-06 | VULM01015833.1 | 1009909 | 1010329 |
| *(predicted orf)* | 3115 bp downstream of TES | -0.10 | 10 | 5.36E-04 | 0.37 | 0.47 | 9.02E-13 | 6.25E-07 | VULM01015885.1 | 1035329 | 1035487 |
| *(predicted orf)* | 3916 bp downstream of TES | -0.11 | 10 | 1.82E-07 | 0.25 | 0.36 | 4.56E-14 | 3.33E-11 | VULM01010276.1 | 2702673 | 2703250 |
| *(predicted orf)* | 4084 bp downstream of TES | -0.15 | 10 | 8.85E-05 | 0.39 | 0.54 | 1.04E-13 | 6.06E-08 | VULM01008304.1 | 1494754 | 1494863 |

### Supplementary Table S5

| -log2(FDR p-value) | FDR p-value | Pathway | DMRs | Total | Genes associated with DMR |
|---|---|---|---|---|---|
| **12.95** | **0.00013** | **Signal transduction** | 52 | (2597) | ARFGAP2, ARHGAP25, ARHGEF7, ARHGEF9, ARL4C, ATP2A3, BUB1, CAVIN1, CCT2, COL27A1, DHRS3, DNAL4, DOCK11, EIF4EBP1, FAM83D, FARP2, FGD3, FGFR3, FRS2, GFRA2, GRM4, HRH3, KSR2, LAMB3, MAMLD1, MAPKAP1, MDM2, MFNG, MYO9A, NCOR2, NR3C1, NRG1, NRTN, PAG1, PARD6A, PDE7B, PDE8B, PDGFRA, PPP1R12B, SCFD1, SH3GL1, SH3KBP1, STRN, TBL1X, TLE4, TNS4, TRHR, USP7, VWF, WNT5B, WWOX, ZNF512B |
| **11.79** | **0.00028** | **MAPK family signaling cascades** | 18 | (588) | EML4, MDM2, STRN, NCOR2, FGFR3, TBL1X, FIP1L1, MAPKAP1, NRG1, GFRA2, NRTN, FRS2, PDGFRA, LMNA, MAMLD1, VWF, LMO7, KSR2 |
| | | *Diseases of signal transduction by growth factor receptors and second messengers* | *16* | *(458)* | *EML4, MDM2, STRN, NCOR2, FGFR3, TBL1X, FIP1L1, MAPKAP1, NRG1, FRS2, PDGFRA, LMNA, MAMLD1, VWF, LMO7, KSR2* |
| | | *MAPK1/MAPK3 signaling* | *8* | *(286)* | *FGFR3, NRG1, GFRA2, NRTN, FRS2, PDGFRA, VWF, KSR2* |
| | | *RAF/MAP kinase cascade* | *8* | *(280)* | *FGFR3, NRG1, GFRA2, NRTN, FRS2, PDGFRA, VWF, KSR2* |
| | | *MAPK family signaling cascades* | *8* | *(325)* | *FGFR3, NRG1, GFRA2, NRTN, FRS2, PDGFRA, VWF, KSR2* |
| **10.35** | **0.00077** | **Signaling by EGFR** | 5 | (59) | SH3GL1, SH3KBP1, FAM83D, PAG1, ARHGEF7 |
| | | *Signaling by EGFR* | *5* | *(53)* | *SH3GL1, SH3KBP1, FAM83D, PAG1, ARHGEF7* |
| | | *EGFR downregulation* | *3* | *(31)* | *SH3GL1, SH3KBP1, ARHGEF7* |
| | | *GAB1 signalosome* | *1* | *(17)* | *PAG1* |
| | | *Negative regulation of MET activity* | *2* | *(21)* | *SH3GL1, SH3KBP1* |
| **9.72** | **0.00119** | **Notch signaling** | 4 | (38) | NCOR2, TBL1X, MAMLD1, MFNG |
| **9.60** | **0.00129** | **PI5P, PP2A and IER3 regulate PI3K/AKT signaling** | 7 | (135) | MDM2, STRN, FGFR3, MAPKAP1, NRG1, FRS2, PDGFRA |
| | | *Negative regulation of the PI3K/AKT network* | *5* | *(113)* | *STRN, FGFR3, NRG1, FRS2, PDGFRA* |
| | | *PI5P, PP2A and IER3 regulate PI3K/AKT signaling* | *5* | *(106)* | *STRN, FGFR3, NRG1, FRS2, PDGFRA* |
| | | *PI3K/AKT signaling in cancer* | *7* | *(104)* | *MDM2, STRN, FGFR3, MAPKAP1, NRG1, FRS2, PDGFRA* |
| | | *Constitutive signaling by aberrant PI3K in cancer* | *5* | *(78)* | *STRN, FGFR3, NRG1, FRS2, PDGFRA* |
| **9.08** | **0.00185** | **Signaling by PDGFR in disease** | 3 | (20) | STRN, FIP1L1, PDGFRA |
| | | *Signaling by PDGFRA extracellular domain mutants* | *1* | *(12)* | *PDGFRA* |
| | | *Signaling by PDGFRA transmembrane, juxtamembrane and kinase domain mutants* | *1* | *(12)* | *PDGFRA* |
| | | *Signaling by cytosolic PDGFRA and PDGFRB fusion proteins* | *2* | *(3)* | *STRN, FIP1L1* |
| | | *Signaling by PDGFR in disease* | *3* | *(20)* | *STRN, FIP1L1, PDGFRA* |
| **8.98** | **0.00198** | **Signaling by receptor tyrosine kinases** | 15 | (535) | COL27A1, FGFR3, MAPKAP1, WWOX, NRG1, SH3GL1, SH3KBP1, DNAL4, TNS4, FAM83D, FRS2, PDGFRA, PAG1, ARHGEF7, LAMB3 |
| **8.00** | **0.00391** | **Circadian clock** | 5 | (86) | NR3C1, TBL1X, HMOX1, CRTC1, CRY1 |
| | | *BMAL1:CLOCK, NPAS2 activates circadian gene expression* | *1* | *(27)* | *TBL1X* |
| | | *Circadian clock* | *4* | *(70)* | *NR3C1, TBL1X, CRTC1, CRY1* |
| | | *RORA activates gene expression* | *1* | *(1)* | *TBL1X* |
| | | *Heme signaling* | *3* | *(48)* | *TBL1X, HMOX1, CRTC1* |
| **7.94** | **0.00407** | **Transmission across chemical synapses** | 12 | (410) | SLC6A13, MDM2, GLUL, NRG1, NRXN1, KCNA3, KCNH5, GRIK3, GRIK4, CAMK1, ARHGEF7, ARHGEF9 |
| | | *Neurotransmitter receptors and postsynaptic signal transmission* | *7* | *(205)* | *MDM2, NRG1, GRIK3, GRIK4, CAMK1, ARHGEF7, ARHGEF9* |
| | | *Transmission across chemical synapses* | *9* | *(270)* | *SLC6A13, MDM2, GLUL, NRG1, GRIK3, GRIK4, CAMK1, ARHGEF7, ARHGEF9* |
| | | *Neuronal system* | *12* | *(410)* | *SLC6A13, MDM2, GLUL, NRG1, NRXN1, KCNA3, KCNH5, GRIK3, GRIK4, CAMK1, ARHGEF7, ARHGEF9* |
| **7.91** | **0.00416** | **RHOU GTPase cycle** | 4 | (54) | MYO9A, ZNF512B, PARD6A, ARHGEF7 |
| | | *RHOV GTPase cycle* | *4* | *(38)* | *MYO9A, ZNF512B, PARD6A, ARHGEF7* |
| | | *RHOU GTPase cycle* | *2* | *(40)* | *PARD6A, ARHGEF7* |
| **7.67** | **0.00491** | **Signaling by ALK in cancer** | 5 | (91) | EML4, MDM2, STRN, FRS2, LMO7 |
| | | *ALK mumants bind TKIs* | *2* | *(12)* | *EML4, STRN* |
| | | *Signaling by ALK fusions and activated protein mutatns* | *5* | *(91)* | *EML4, MDM2, STRN, FRS2, LMO7* |
| | | *Signaling by ALK in cancer* | *5* | *(91)* | *EML4, MDM2, STRN, FRS2, LMO7* |
